# Neuronal and glial transcriptional states vary with proximity to tangle-bearing neurons in the human entorhinal cortex

**DOI:** 10.64898/2026.10.27.762234

**Authors:** Samuel L. Boulger, Nathalie Lambie, Jan Wos, Marianna Papageorgopoulou, Sabino Méndez Pastor, Emily Adair, Michael Thomas, Alessia Caramello, Nurun N. Fancy, Johanna S. Jackson, Eugene P. Duff, Anna Mallach, Paul M. Matthews

## Abstract

More neurons die in Alzheimer’s disease than form tangles, and tangle-bearing neurons can persist for decades, so tangles alone cannot explain degeneration. We combined 6,000-plex spatial transcriptomics with PHF1 immunofluorescence (167,176 cells; 9 donors) and imaging mass cytometry (170,195 cells; 62 donors), measuring each cell’s distance to the nearest tangle. Tangle-bearing layer 2-3 excitatory neurons were enriched late along a trajectory of increasing microtubule-associated, synaptic and V-ATPase expression, with elevated p62 and K48-ubiquitin. Tangle-free neurons shared more of this state the closer they lay to a tangle, including higher chaperone and autophagy scores, p62 and nuclear size. Nearby glia had higher disease- associated scores, with astrocytes expressing less GFAP. Excitatory, but not inhibitory, neuron loss was greatest in subtypes closest to tangles and cortical depths richest in phosphorylated tau. Selective neuronal vulnerability may reflect not only a neuron’s susceptibility to forming a tangle but also where it lies relative to one.

**Highlights:**

- Single-cell spatial analysis defines a tangle microenvironment in human cortex
- Tangle-free neurons near tangles have higher stress and autophagy gene expression
- Microglia and astrocytes near tangles show disease-associated reactive signatures
- Excitatory neuron loss is concentrated where local phosphorylated tau is highest

## Introduction

Neurofibrillary tangles are somatodendritic inclusions of paired helical and straight filaments^1^ assembled from the microtubule-associated protein tau. Hyperphosphorylation reduces the affinity of tau for microtubules^2^ and expands the free cytosolic pool, promoting redistribution of tau from the axon to the soma and dendrites^3^ where the microtubule-binding repeats adopt the ordered cross-β fold that defines Alzheimer’s disease (AD) pathology^4,5^.

Tau phosphorylation is not intrinsically pathological. Hibernating mammals accumulate tau phosphorylated at PHF-like epitopes during torpor and reverse this after arousal without filament formation^6,7^, and hypothermia induced by anaesthesia produces hyperphosphorylation through inhibition of protein phosphatase 2A (PP2A) that is rescued by restoring normal body temperature^8^. Fetal tau is phosphorylated to an extent comparable to PHF-tau and at shared sites, declining with maturation^9,10^. Tau is also required for hippocampal long-term depression, the induction of which is associated with glycogen synthase kinase 3 (GSK3)-mediated tau phosphorylation^1^^1^, and site-specific phosphorylation at Thr205 protects against Aβ-induced excitotoxicity^12^. What distinguishes AD is therefore not the presence of phosphorylated tau but the reduction in its reversibility. PP2A, which accounts for most tau phosphatase activity in the human brain, is reduced in activity, expression and methylation^13–15^, while cyclin-dependent kinase 5 (CDK5) activity rises through accumulation of its truncated activator p25^16^. GSK3 accumulates in tangle-bearing neurons without increased activity, suggesting that, with reduced phosphatase activity, even normal GSK3 activity may suffice to shift the equilibrium towards phosphorylation^17^. Once tau assembles into filaments, it leaves this equilibrium altogether, with part of its microtubule-binding repeat region locked into an ordered cross-β core^5^.

The relationship between tangle formation and neuronal death is less straightforward than their general association implies. At the regional level, tangle burden tracks clinical severity^18^; however, at the level of the individual neuron, the presence of a tangle does not identify a cell at elevated risk of death. Stereological counts reveal that neuronal loss in AD exceeds the number of tangles present^19,20^, and longitudinal multiphoton imaging in mouse tauopathy models shows that tangle-bearing neurons persist for months *in vivo* without dying^21^, retain stable resting calcium levels and are functionally responsive^22^. One study found that tangle-bearing neurons die at less than a third of the rate of their tangle-free counterparts^23^. Modelling of human post-mortem counts likewise estimates tangle-bearing neuron survival on the order of decades^24^. *In vivo*, tangle-bearing neurons with activated executioner caspases do not acutely degenerate^25^, and caspase activation precedes filament assembly in neurons that go on to form a tangle and survive^21^.

If the neurons that die in AD and other tauopathies are not, for the most part, the neurons that bear tangles, then the pathological consequences associated with tangle burden are not accounted for by the tangle-bearing cell alone. Two possibilities follow. A tangle-bearing neuron may be altered in ways that compromise its function and that of its circuit without killing it, or it may harm the surrounding tissue, directly through release of soluble tau and loss of synaptic input, or indirectly through altered states in local glia. These are not mutually exclusive, and neither can be addressed by approaches that isolate cells from their tissue context. Single-soma transcriptomic profiling of AT8-positive neurons^26^ and laser-capture proteomics of tangle-bearing cells^27^ have characterised the state of tangle-bearing cells with considerable precision, while discarding the spatial information needed to ask what is happening to the cells nearby.

We therefore combined spatial transcriptomics of human entorhinal cortex (EC) from donors spanning a range of AD neuropathological change with imaging mass cytometry (IMC) of the same region in a larger cohort reported separately^28^. Phosphorylated tau (PHF1, pSer396/pSer404) was imaged on the profiled sections in each modality – by immunofluorescence for spatial transcriptomics and as a channel in IMC – allowing tangle status to be assigned to individual segmented cells and enabling every cell to be analysed in terms of proximity to the nearest tangle-bearing neuron within its own tissue section. We asked three questions. First, how does the transcriptome of a tangle-bearing neuron differ from that of a neuron of the same subtype without a tangle? Second, do tangle-free neurons near a tangle-bearing neuron differ from tangle-free neurons further away? Third, do microglia and astrocytes near tangle-bearing neurons differ in state from those further away?

## Results

### Spatially resolved single-cell transcriptomic profiling of the human EC

Spatial transcriptomics data were acquired for EC from 9 human donors with entorhinal tau pathology spanning Braak stages I, IV and VI using the CosMx Human 6K Discovery Panel (Figure 1A; Table S1). Cells were first annotated into broad classes using marker gene-based annotation from a single-nucleus RNA sequencing EC reference^28^ (Figure 1B; Table S2). Neurons underwent a second round of marker-based annotation to produce more granular classes based on gene lists from the same reference (Figure 1C). Cell class proportions were consistent with published estimates for EC^29^ (Figure 1D). Low-quality cells were excluded (see STAR Methods), leaving 167,176 cells across 9 donors (Table S3). After CosMx imaging, the same sections were immunostained for PHF1 (phospho-tau pSer396/pSer404) to identify neurons bearing somatic phospho-tau inclusions. PHF1 binds phosphorylated tau irrespective of assembly state, labelling pre-tangle somatic tau, neuropil threads and dystrophic neurites as well as inclusions, so per-cell intensity alone may not accurately identify the presence of an inclusion. Inclusions were therefore identified by manual annotation of somatic morphology and classified as neurofibrillary tangles on morphological criteria, with final annotations made after review by three raters (Figure 1E, see STAR Methods). Blinded re-annotation of 600 neurons (150 tangle-bearing, 450 tangle-free) gave 94.8% agreement (κ = 0.86, 95% CI 0.77– 0.91; AC1 = 0.92, 0.88–0.95), corresponding to strong intra-rater agreement^30^. Manual annotations were compared against normalised per-cell PHF1 intensity as a researcher- independent quantitative measure (Figure 1F). Tangle-bearing neurons had a median intensity 7.9 robust s.d. above their sample median (IQR 5.6–11.0) and intensity discriminated manual calls with ROC AUC 0.982 (range 0.93–1.00 across donors; Figure 1F). Based on the manual annotations, neurons in the CosMx data are described as “tangle-bearing” and “tangle-free”, while in the IMC data neurons are described as PHF1_high_ and PHF1_low_.

**Figure 1.**
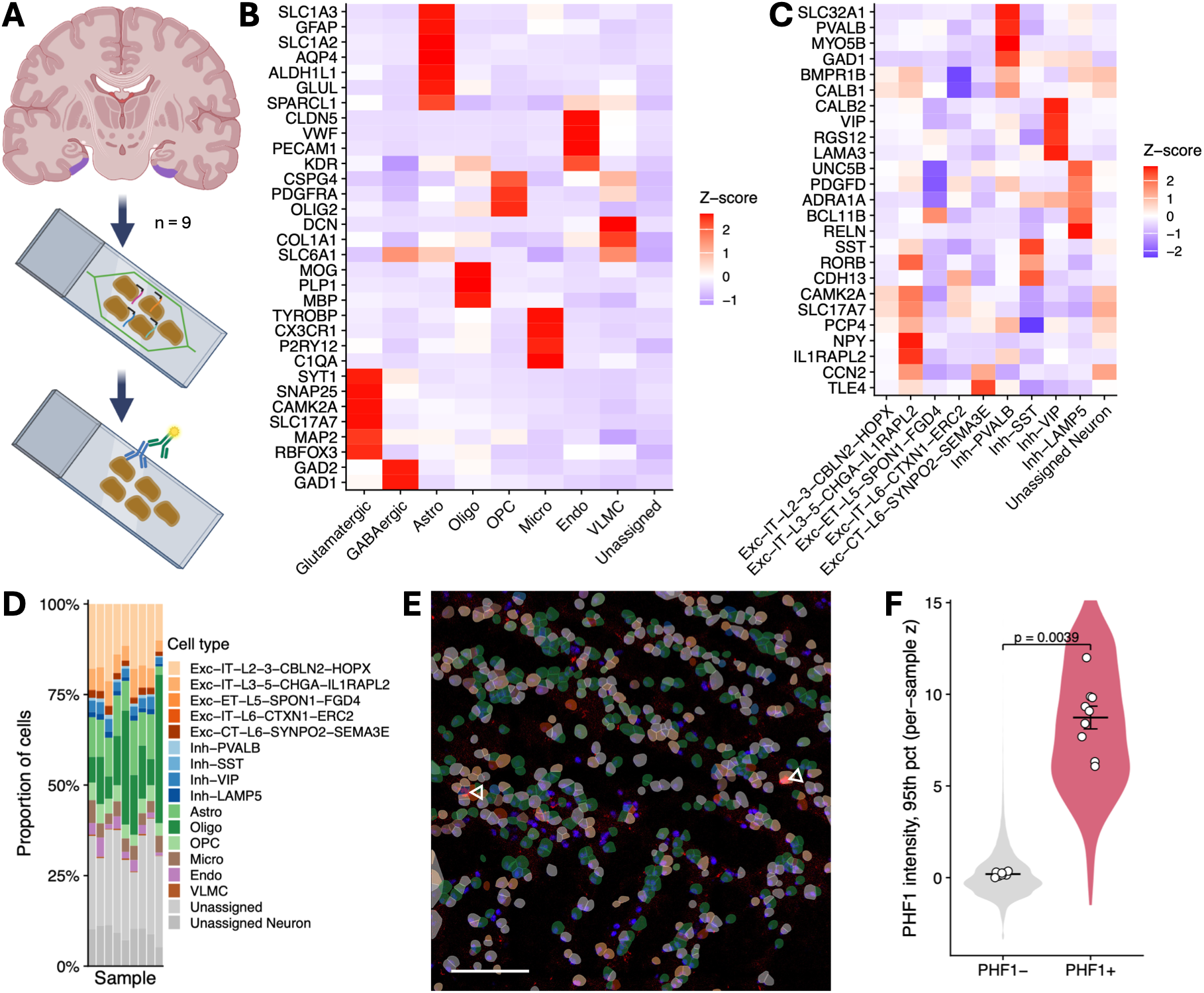
Paired 6,000-plex spatial transcriptomics and PHF1 immunofluorescence in human EC. **(A)** EC tissue from 9 donors (three each at Braak stages I, IV and VI) was profiled by CosMx spatial molecular imaging with post-run PHF1 phospho-tau immunofluorescence. **(B and C)** Mean expression (per-gene z-score) of canonical marker genes across broad cell classes (**B**) and neuronal subtypes **(C)**, each predicted by sequential marker gene-based annotation from an entorhinal cortex snRNA-seq reference^28^. **(D)** Per-donor cell-type composition. Neurons represented 37.8% of all cells (donor-level 95% CI 31.7–44.0%). This is comparable to the 36.6% reported in a recent single-cell atlas^29^. **(E)** Representative segmented CosMx field of view showing DAPI (blue) and PHF1 stain (red), with cell segmentation coloured by cell type as in **D**; white arrowheads mark manually annotated tangle-bearing cells. Scale bar, 100 μm. **(F)** Per-cell PHF1 95^th^ percentile mask intensity recovers manual tangle calls. Violins show the cell-level distribution of per-sample *z*-scored PHF1 intensity for tangle-free (PHF1-) and tangle-bearing (PHF1+) neurons pooled across donors; points are per-donor means, horizontal bars and whiskers denote mean ± s.e.m. across donors. *p* = 0.0039, two-sided paired Wilcoxon signed-rank test on donor means (*n* = 9).

### Tangle-bearing excitatory neurons are enriched at the end of a transcriptional trajectory marked by microtubule-associated and synaptic genes

We first asked what distinguishes a neuron bearing a neurofibrillary tangle from other neurons of the same subtype. We focused on the Exc-IT-L2-3-*CBLN2*-*HOPX* neuronal population, which had the highest density of tangle-bearing neurons with a mean of 1.42 per mm^2^ (95% CI 0.51–2.34; Figure 2A). Because tangle-bearing neurons are rare (188 of 26,585 in this subtype), we ordered all neurons of the subtype along a tangle-associated pseudotime trajectory, as done previously^27^. Tangle-bearing neurons were situated further along the trajectory than tangle-free neurons from the same donor (Figure 2B), and their proportion of the population increased along the trajectory (Figure 2C). The tangle-associated transcriptional trajectory recapitulated the tangle-associated signature previously described^26^ in layer 2-3 neurons. The set of genes reported to be upregulated with AT8 positivity in these neurons scored higher in tangle-bearing than tangle-free neurons in all tested donors and its score increased along the trajectory (Figures 2D and 2E). By testing each gene against pseudotime with a linear mixed model, we found that expression of 87 genes increased and expression of 295 genes decreased by more than 0.25 log_2_ units across the tangle-associated trajectory (adjusted *p* < 0.1; Table S4).

**Figure 2.**
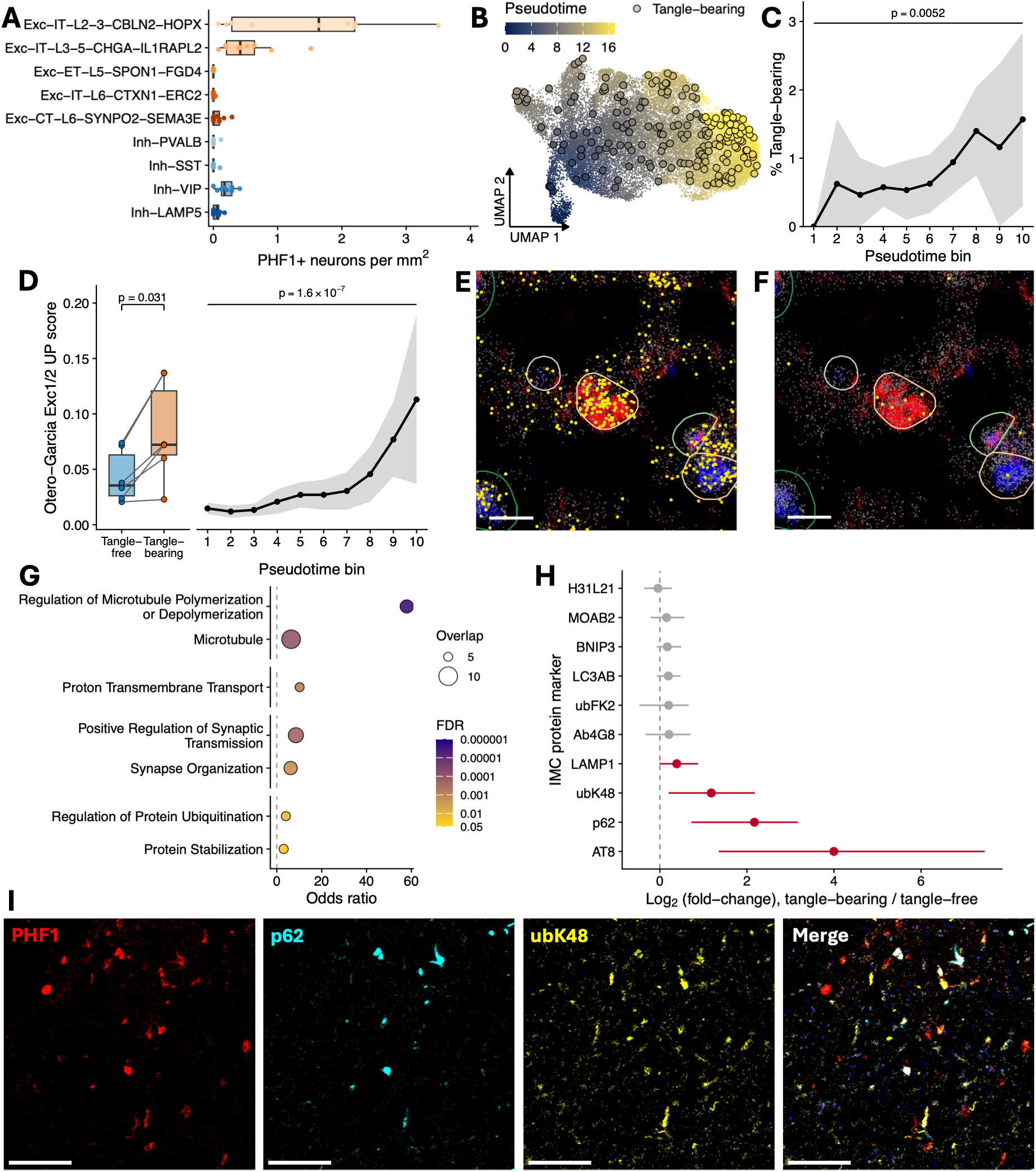
Tangle-bearing neurons are enriched towards the end of a transcriptional trajectory that reproduces published tangle signatures. **(A)** Number of manually annotated tangle-bearing neurons per *mm^2^* by neuronal subtype. Exc-IT-L2-3-*CBLN2*-*HOPX* had the highest density with a mean of 1.42 tangle-bearing cells per *mm^2^*(95% CI 0.51–2.34) and a raw count mean of 20.9 tangle-bearing neurons per donor (6.2–35.5; *n* = 9). The density for all subtypes combined was 2.30 tangle-bearing neurons per mm^2^ (1.01–3.59). **(B)** UMAP of Exc-IT-L2-3-*CBLN2*-*HOPX* neurons (*n* = 26,585 cells, 9 donors) coloured by pseudotime. Larger circles, tangle-bearing neurons (*n* = 188). Tangle-bearing neurons were situated further along the trajectory than tangle-free neurons (donor-mean AUC 0.61; empirical *p* = 0.002, within-donor label permutation). **(C)** Percentage of Exc-IT-L2-3-*CBLN2*-*HOPX* neurons that are tangle-bearing in ten equal-width pseudotime bins. Line, mean of per-donor percentages; shading, 95% CI (clipped at 0). *n* = 6 donors with ≥10 tangle-bearing neurons (20,866 neurons, 182 tangle-bearing). The tangle-bearing percentage differed across bins (Friedman test, χ^2^(9) = 23.5, *p* = 0.0052, Kendall’s W = 0.44). **(D)** Gene set score of the previously reported Exc1/2 tangle-associated upregulated signature^26^ (Seurat AddModuleScore, 319 genes) in the donors and neurons of (**C**). Left, mean score per donor in tangle-free and tangle-bearing neurons; lines join each donor’s pair; box, median and interquartile range; whiskers, 1.5× interquartile range. The score was higher in tangle-bearing neurons (mean paired difference 0.040, 95% CI 0.015–0.065; Wilcoxon signed-rank test, V = 21, *p* = 0.031). Right, score across the pseudotime bins of (**C**) in all neurons; line, donor mean; shading, 95% CI. The score differed across bins (Friedman test, χ^2^(9) = 49.1, *p* = 1.6 × 10^-7^, Kendall’s W = 0.91). *n* = 6 donors with ≥10 tangle-bearing neurons each. **(E and F)** Representative images centred on a manually annotated tangle-bearing (red) Exc-IT-L2-3-*CBLN2*-*HOPX* neuron, with DAPI in blue, all transcripts in grey, and transcripts from the previously reported Exc1/2 tangle-associated upregulated gene list^26^ (**E**) or transcripts of genes in the overrepresented regulation of microtubule polymerization or depolymerization term of **G** (**F**) in yellow. Cell outlines represent the segmented cell masks and are coloured by cell type as in **A** and Figure 1D. Scale bar, 10 μm. **(G)** Enriched GO terms among genes with increasing expression along the Exc-IT-L2-3-*CBLN2*-*HOPX* tangle-associated trajectory (linear mixed model with donor random intercept; FDR < 0.1, |log_2_FC| > 0.25). *x*-axis, odds ratio; point size, number of overlapping genes; fill, BH-adjusted *p*. Terms shown were selected for comparison with previously reported tangle- associated categories^26,27^ (7 of 34 enriched terms); the full list is in Table S5. **(H)** Differential protein abundance in PHF1_high_ versus PHF1_low_ excitatory neurons in published imaging mass cytometry (IMC) data^28^ (*n* = 36,133 neurons from 44 AD donors). Ten cell-state markers were tested. Points, model estimate (PHF1_high_ vs PHF1_low_, adjusted for excitatory subcluster); bars, 2.5^th^–97.5^th^ percentiles of donor differences (n = 44). Per-marker linear mixed models (marker ∼ PHF1 status + subcluster + sex + age + PMI + (1|donor)); red, BH-adjusted *p* < 0.05 and |log_2_FC| ≥ 0.25; grey, all others. **(I)** Representative 400 × 400 μm layer 2-3 subregion of an IMC image. Red, PHF1; cyan, p62; yellow, ubK48; navy, DNA. Scale bar, 100 μm.

Genes with increasing expression along the tangle-associated trajectory were enriched for terms in categories previously reported in tangle-bearing neurons^26,27^, including regulation of microtubule polymerisation, driven by *MAPT*, *MAP1B*, and the stathmins *STMN1-4*, and for synaptic transmission and organisation pathways (Figures 2F and 2G; Table S5). Genes with higher expression along the trajectory also included V-type ATPase subunits responsible for endolysosomal acidification via proton transmembrane transport (*ATP6V1B2*, *ATP6V1E1*, *ATP6V0C*) and genes involved in the regulation of protein ubiquitination (*UBB*, *HSP90AA1*, *HSP90AB1*, *DNAJA1*).

To determine whether these transcriptional differences related to proteostasis pathways extend to the protein level, we contrasted immunostaining intensity of 10 disease-associated proteins, including several related to protein clearance, between PHF1_high_ and PHF1_low_ excitatory neurons in the 44 donors with AD from a multiplexed imaging dataset generated previously^28^. AT8, which recognises tau phosphorylated at Ser202 and Thr205, served as a positive control and was elevated in PHF1_high_ excitatory neurons (44 of 44 donors), as were LAMP1 (42 of 44), the autophagy receptor p62 and K48-linked polyubiquitin (each 44 of 44; Figures 2H and 2I). K48-linked conjugates were elevated more than total ubiquitin conjugates detected by FK2 (log_2_FC 1.16 vs 0.24), consistent with buildup of proteasome-targeted substrates in PHF1_high_ neurons.

### Elevated stress- and autophagy-related gene expression in tangle-free neurons close to tangle-bearing neurons

The distance to the nearest tangle-bearing neuron was calculated for every tangle-free cell in the CosMx dataset to investigate transcriptional differences associated with proximity to tangle-bearing neurons. We first measured PHF1 signal intensity in tangle-free Exc-IT-L2-3-*CBLN2*-*HOPX* neurons to investigate the spatial distribution of subthreshold tau phosphorylation in relation to tangle-bearing neurons. Tangle-free neurons situated nearer tangle-bearing neurons had greater PHF1 signal intensity than tangle-free neurons situated further away (Figure 3A), indicating a gradient of subthreshold PHF1 immunoreactivity within the cell masks of neurons surrounding tangle-bearing neurons.

**Figure 3.**
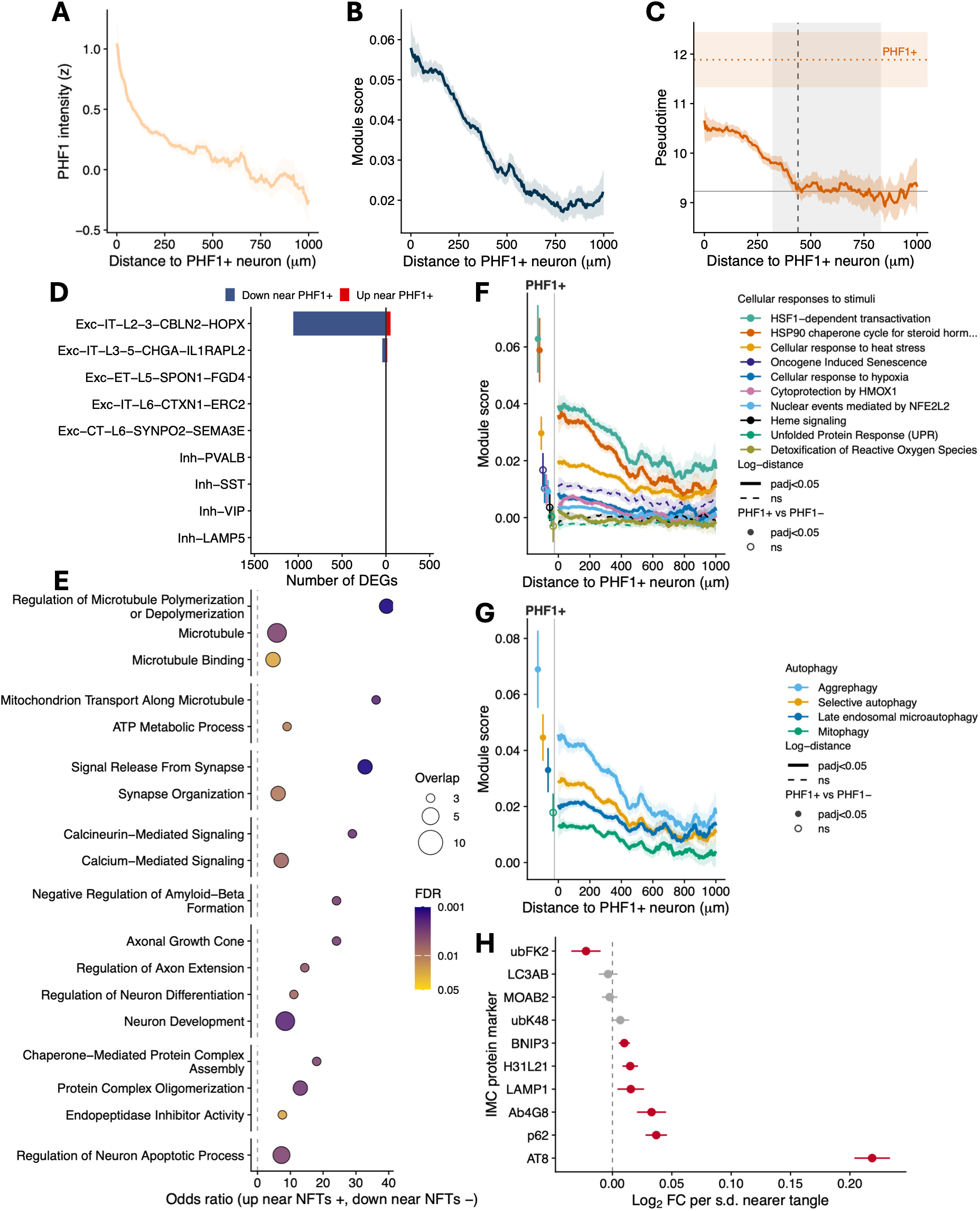
Transcriptional state in tangle-free excitatory neurons varies with distance to the nearest tangle-bearing neuron. **(A)** PHF1 intensity (per-cell 95^th^ percentile, robust *z* within sample) in tangle-free Exc-IT-L2-3-*CBLN2*-*HOPX* neurons versus distance to the nearest tangle-bearing neuron; line, 75 μm unadjusted rolling mean; shading, 95% CI. Linear mixed model on log-distance (sex, age, PMI, donor random intercept): β = -0.26 per s.d. log-distance (s.d. = 0.77 log-units; 95% CI -0.28, -0.24); *p* = 9.1 × 10^-152^. *n* = 24,657 neurons, 9 donors. **(B)** Per-cell module scores in tangle-free Exc-IT-L2-3-*CBLN2*-*HOPX* neurons over distance to the nearest tangle-bearing neuron, for a previously reported phospho-tau-associated gene set^26^. Lines, 75 μm rolling mean after regressing out per-cell PHF1 intensity (95^th^ percentile within the cell mask, robust *z* within sample); shading, 95% CI (24,657 cells, 9 donors). Linear mixed model on log-distance, donor random intercept, adjusted for total counts, negative probe fraction, sex, age and PMI (β = -0.0034, *p* = 1.1 × 10^-21^). Distance and intensity were only weakly correlated (*r* = -0.15, variance inflation factor 1.03), and intensity uniquely explained 0.12% of module-score variance in tangle-free neurons (semipartial *R*^2^). Adjustment steepened the slope relative to unadjusted fits (β = -0.0029), and the slope was unchanged with intensity entered as a within-donor rank, natural spline (df = 3) or top decile indicator and persisted within the lowest intensity decile alone (β = -0.0040, *p* = 1.9 × 10⁻^4^). **(C)** Tangle-associated pseudotime scores (Figure 2B) in tangle-free Exc-IT-L2-3-*CBLN2*-*HOPX* neurons over distance from nearest tangle-bearing neuron. Pseudotime increased with proximity to tangle-bearing neurons (linear mixed model, -0.27 pseudotime units per s.d. log-distance, 95% CI -0.32 to -0.21, *p* = 6.9 × 10^-21^, *n* = 24,672 neurons from 9 donors). Continuous orange line, 75 μm rolling mean; dotted orange line, mean pseudotime value in manually annotated tangle-bearing neurons; orange shaded regions, 95% CI; dashed grey line, breakpoint of a hinge model on raw distance (flat beyond the breakpoint; donor fixed effects, nUMI and negative-probe covariates; grey shaded area, donor-bootstrap 95% CI). Pseudotime was higher than its distal baseline within 440 μm of a tangle-bearing neuron (95% CI 320–830 μm). **(D)** Number of significant differentially expressed genes at BH-adjusted *p* < 0.1 per neuronal subtype. Distance-dependent differential expression among tangle-free neurons within 1,000 μm of a tangle-bearing neuron (per-gene linear mixed model, variancePartition dream on log-CPM versus s.d.-scaled natural-log distance: expression ∼ log-distance + nUMI + negative-probe fraction + sex + age + PMI + (1|donor); BH-corrected). **(E)** Genes at BH-adjusted *p* < 0.1 in Exc-IT-L2-3-*CBLN2*-*HOPX* neurons were tested for GO overrepresentation (enrichR; per-cell-type expressed background; genes with increasing and decreasing expression separately; terms at BH-adjusted *p* ≤ 0.05). Direction is shown in proximity terms (higher near tangles, positive). *x*-axis, proximity-signed odds ratio; point size, overlapping DEGs; fill, BH-adjusted *p*; terms grouped by function. **(F and G)** Gene sets from the ‘cellular responses to stimuli’ (**F**) and ‘autophagy’ (**G**) Reactome clusters were filtered (see STAR Methods) and scored (AddModuleScore) in tangle-free Exc-IT-L2-3-*CBLN2*-*HOPX* neurons against distance to the nearest tangle-bearing neuron (*n* = 24,672 cells, 9 donors) and in tangle-bearing versus tangle-free Exc-IT-L2-3-*CBLN2*-*HOPX* neurons (*n* = 26,585 neurons, 188 tangle-bearing, 9 donors). Lines, unadjusted rolling means (75 μm window) ± 95% CI; solid lines, significant log-distance term in a linear mixed model as in **B**; dashed lines, adjusted *p* > 0.05. β is the difference in module score per s.d. log-distance, so a negative value denotes a higher score near tangles. s.d. = 0.77 log-units. Scores (β [95% CI]) were elevated near tangles for cellular response to heat stress (-0.00069 [-0.0011, -0.00031]; adjusted *p* = 0.00097), nuclear events mediated by NFE2L2 (-0.00052 [-0.00083, -0.00021]; adjusted *p* = 0.0018), cellular response to hypoxia (-0.00070 [-0.0011, -0.00031]; adjusted *p* = 0.00097), HSP90 chaperone cycle for steroid hormone receptors (SHR) in the presence of ligand (-0.0021 [-0.0028, -0.0013]; adjusted *p* = 1.8 × 10^-7^), cytoprotection by *HMOX1* (-0.00060 [-0.0010, -0.00018]; adjusted *p* = 0.0071), detoxification of reactive oxygen species (-0.00072 [-0.0012, -0.00026]; adjusted *p* = 0.0036), and HSF1-dependent transactivation (-0.0016 [-0.0024, -0.00086]; adjusted *p* = 0.00015) in **F**, and for selective autophagy (-0.0013 [-0.0019, -0.00083]; adjusted *p* = 5.8 × 10^-7^), aggrephagy (-0.0023 [-0.0031, -0.0015]; adjusted *p* = 1.2 × 10^-7^), mitophagy (-0.00068 [-0.0012, -0.00014]; adjusted *p* = 0.016), and late endosomal microautophagy (-0.00072 [-0.0013, -0.00014]; adjusted *p* = 0.016) in **G**. Marginal points at left show mean ± 95% CI for tangle-bearing neurons; filled points, significant tangle-bearing versus tangle-free contrast from the equivalent model; open points, non-significant. Tangle-bearing neurons showed higher cellular response to heat stress (adjusted *p* = 0.00067, donor-paired *d_z_* = 0.54, 7/9 donors concordant), nuclear events mediated by NFE2L2 (adjusted *p* = 0.0042, *d_z_* = 0.72, 6/9), HSP90 chaperone cycle for SHR (adjusted *p* = 7.7 × 10^-6^, *d_z_* = 0.45, 6/9), and HSF1-dependent transactivation (adjusted *p* = 6.3 × 10^-5^, *d_z_* = 0.54, 7/9) scores in **F**, and higher selective autophagy (adjusted *p* = 7.3 × 10^-5^, *d_z_* = 0.60, 6/9), aggrephagy (adjusted *p* = 3.1 × 10^-5^, *d_z_* = 0.38, 6/9) and late endosomal microautophagy (adjusted *p* = 0.0084, *d_z_* = 0.71, 7/9) scores in **G**. All eleven gradients retained their sign in every leave-one-donor-out refit; nine also retained significance in all nine folds (cytoprotection by *HMOX1* in 8, mitophagy in 7). **(H)** Protein gradients around PHF1_high_ neurons in EC excitatory neurons. Log_2_FC per s.d. decrease in log-distance to the nearest PHF1_high_ neuron (IMC; 26,653 PHF1_low_ excitatory neurons, 44 AD donors, ≤300 μm); positive values indicate higher intensity near PHF1_high_ neurons. Per-marker linear mixed models (marker ∼ distance + subcluster + sex + age + PMI + (1|donor)); points, estimate; bars, 95% CI; red, BH-adjusted *p* < 0.05; grey, not significant.

We next scored expression, over distance to the nearest tangle-bearing neuron, of the set of genes previously reported as upregulated in AT8-positive neurons^26^ and shown to score higher along our tangle-associated trajectory (Figure 2D). The module score increased with proximity to tangle-bearing neurons in tangle-free Exc-IT-L2-3-*CBLN2*-*HOPX* neurons (Figure 3B), indicating that these neurons share more of the differentially expressed, tangle-associated transcriptional state the closer they lie to a tangle-bearing neuron. Adjustment for per-cell PHF1 intensity did not attenuate the gradient (Figure S1A; β per s.d. log-distance = −2.9 × 10^-3^ unadjusted, *p* = 5.1 × 10^-17^, versus −3.4 × 10^-3^ adjusted, *p* = 1.1 × 10^-21^). The gradient also persisted with the exclusion of neurons within 200 μm of a tangle and with PHF1 intensity decile stratification (Figures S1B and S1C), indicating that the gradients among tangle-free neurons are not carried by cells immediately adjacent to tangle-bearing neurons or by the neuron’s own subthreshold PHF1 signal.

Module scores can shift with technical factors such as transcript detection or cell quality, which could themselves vary with distance. We therefore scored a negative-control module of 305 genes stably expressed in this subtype in our snRNA-seq reference^28^ (scSEGIndex). Its slope was statistically significant but negligible in size (β = -3.6 × 10^-4^ per s.d. log-distance; 95% CI -5.3 × 10^-4^ to -1.9 × 10^-4^; adjusted *p* = 2.1 × 10^-4^; Figure S2A). This module therefore provides an empirical floor for non-specific distance-dependent module-scoring, which the reported gradient (Figure 3B) exceeds 9-fold.

Proximity likewise tracked position along the tangle-associated trajectory. Tangle-free Exc-IT-L2-3-*CBLN2*-*HOPX* neurons situated closer to tangle-bearing neurons were further along the tangle-associated transcriptional trajectory than more distant neurons (Figure 3C), indicating increasing transcriptional similarity to tangle-bearing neurons with increased proximity. We next modelled gene expression in tangle-free neurons as a function of distance to the nearest tangle-bearing neuron. Exc-IT-L2-3-*CBLN2*-*HOPX* (1,110 significant genes; adjusted *p* < 0.1; Table S6) and Exc-IT-L3-5-*CHGA*-*IL1RAPL2* (58) accounted for 98.8% of distance-associated differentially expressed genes in neurons (Figure 3D). In the Exc-IT-L2-3-*CBLN2*-*HOPX* neurons, genes with higher expression closer to tangle-bearing neurons included microtubule-associated (*MAPT*, *MAP1B*, *STMN1*, *STMN3*, *TUBB2A*), 14-3-3 protein (*YWHAG*, *YWHAH*, *YWHAZ*), calmodulin (*CALM1*, *CALM3*), presynaptic (*SNAP25*, *SYT1*, *SYN2*, *CPLX1*) and proteostasis-related genes (*UBB*, *CLU*, *HSP90AA1*, *PRNP*, *APP*), and were enriched for Gene Ontology terms related to microtubule organisation, chaperone-mediated complex assembly, synaptic and calcium signalling, suppression of Aβ formation, apoptosis, and developmental and regenerative pathways (Figure 3E; Table S7).

To test whether proximity effects were driven by transcripts of tangle-bearing neurons being assigned to adjacent cells during segmentation, we refitted the same differential gene expression model to Exc-IT-L2-3-*CBLN2*-*HOPX* neurons after excluding cells within 10, 20, 30, and 50 μm of a tangle-bearing neuron. Gene-level coefficients from the sensitivity tests correlated closely with those from the primary analysis (Spearman ρ = 0.99, 10 μm; 0.96, 20 μm; 0.94, 30 μm; 0.89, 50 μm; all *p* < 10^-300^; Figure S3A; Table S8). To test whether proximity effects reflected cortical depth, we refitted the distance model with scaled cortical depth as an additional covariate. Gene-level distance coefficients correlated closely with those from the primary analysis (ρ = 0.995; *p* < 10^-300^; Figure S3B). The observed proximity effects are therefore not explained by transcripts of tangle-bearing neurons being assigned to adjacent cells during segmentation, or by the cortical depth of the tangle-free neurons. Adding a random intercept for imaging field of view, so that distance effects were estimated only within fields of view, gave similar coefficients (ρ = 0.99; *p* < 10^-300^; Figure S3C). Finally, because a single section cannot detect tangles above or below it, we simulated tangles at the observed density in the unimaged tissue 500 μm either side of each section and recomputed each cell’s distance to the nearest tangle in three dimensions. This reduced the median distance from 266 to 107 μm, with the largest reductions for the most distant cells. Gene-level coefficients still correlated closely with those from the primary analysis (ρ = 0.86; *p* < 10^-300^; Figure S3D). No significant differentially expressed gene from the primary model changed sign in any of the sensitivity tests.

We next asked whether tangle-bearing neurons, or the neurons around them, show transcriptional signatures of stress or cell death. We tested three Reactome subtrees: cellular responses to stimuli, to capture stress responses; autophagy, since phospho-tau is a substrate for degradative clearance; and programmed cell death. Within each subtree, gene sets were filtered on set size, panel coverage and non-redundancy alone, without reference to expression (see STAR Methods). We scored Exc-IT-L2-3-*CBLN2*-*HOPX* neurons for these gene sets and contrasted scores in tangle-bearing versus tangle-free neurons and in tangle-free neurons over distance to the nearest tangle-bearing neuron. We observed higher scores for two intrinsic apoptosis Reactome modules (Figure S4A) and found that this was attributable to increased expression of genes annotated as negative regulators of apoptosis including the *YWHA* family of genes^31^ both in tangle-bearing neurons (Figure S4B) and in tangle-free neurons with increasing proximity to the nearest tangle-bearing neuron (Figure S4C). Tangle-bearing neurons and tangle-free neurons close to tangles both had elevated module scores for HSF1- dependent transactivation, HSP90 chaperone cycle for steroid hormone receptors, cellular response to heat stress, and nuclear events mediated by NFE2L2 (Figure 3F). Scores for the cellular response to hypoxia, reactive oxygen species detoxification, and cytoprotection by *HMOX1* gene expression modules were elevated with proximity but did not differ significantly between tangle-bearing and tangle-free neurons (Figure 3F).

Scores for the Reactome aggrephagy, selective autophagy, and late endosomal microautophagy modules were elevated both in tangle-bearing relative to tangle-free neurons and in tangle-free neurons with increasing proximity to tangle-bearing neurons, whereas the mitophagy module score increased with proximity but did not differ significantly between tangle-bearing and tangle-free neurons (Figure 3G). In the IMC AD cohort, p62 protein was elevated in PHF1_high_ neurons (Figure 2H) and in PHF1_low_ neurons close to PHF1_high_ neurons (Figure 3H). LC3A/B abundance was not elevated in either population. FK2-detected ubiquitin conjugates were lower with proximity to PHF1_high_ neurons, with no significant difference in abundance of K48-linked chains (Figure 3H), in contrast to the co-occurrence of both p62 and K48-linked conjugates in PHF1_high_ neurons themselves (Figure 2H).

### Excitatory neuron loss in AD is concentrated where local phosphorylated tau burden is highest

Selective neuronal vulnerability in AD describes the differential susceptibility of individual neuronal subtypes to degeneration. Using the IMC data^28^, we calculated the percentage difference in columnar density – the number of cells per mm along the cortical ribbon spanning the full grey matter depth – of each neuronal cluster between non-diseased (Braak 0–I; *n* = 18) and AD (Braak III, IV, VI; *n* = 44) donors. We compared this with each cluster’s geometric mean distance to nearest PHF1_high_ neuron in AD. Clusters lying closer to PHF1_high_ neurons were more depleted (Figure 4A; Spearman ρ = 0.66, 95% CI 0.14 to 0.83; permutation *p* = 0.044), most prominently Exc 2 (CALB1). In contrast, cluster-level loss was not significantly associated with PHF1 intensity (Figure 4B; ρ = -0.44, 95% CI -0.73 to 0.05; permutation *p* = 0.21); with only ten clusters, the confidence interval does not exclude a substantial association.

**Figure 4.**
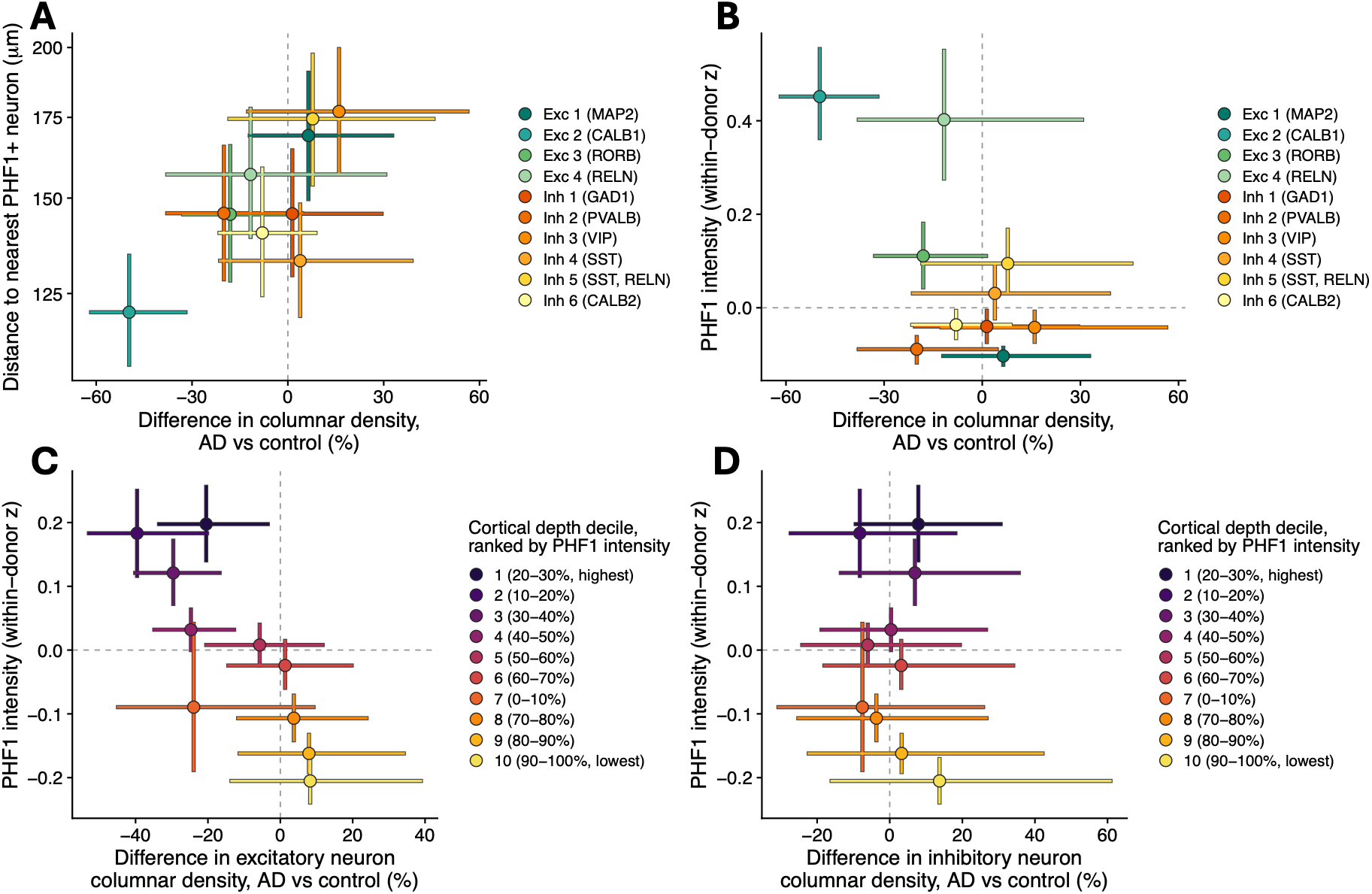
Phosphorylated tau burden tracks the neuronal populations and cortical depths depleted in AD. **(A and B)** Difference in columnar density between non-diseased (Braak 0–I; *n* = 18 donors) and AD (Braak III, IV, VI; *n* = 44 donors) for each of the 10 neuronal clusters, plotted against (**A**) geometric-mean distance from PHF1_low_ cells to the nearest PHF1_high_ neuron, or (**B**) PHF1 intensity (within-donor z-score) in AD. **(C and D)** Each ROI was divided into ten relative cortical-depth bands (deciles, from the pia). Columnar density difference in (**C**) excitatory or (**D**) inhibitory neurons, plotted against PHF1 intensity of all neurons in AD. Bands are ranked by PHF1 intensity (1, highest). Points show cohort-level estimates, and error bars show 95% CIs from resampling donors (AD and control separately). Spearman correlation across the 10 clusters (**A and B**) or 10 depth bands (**C and D**): (**A**) ρ = 0.66, *p* = 0.044; (**B**) ρ = -0.44, *p* = 0.21; (**C**) ρ = -0.82, *p* = 0.006; (**D**) ρ = -0.15, *p* = 0.68 (permutation tests).

Because tangles are unevenly distributed throughout the cortex, we next asked whether neuronal loss co-varies with tangle pathology across cortical depth. We divided every region of interest (ROI) into ten relative-depth bands (deciles, from the pia). For each band, we compared excitatory neuron density between non-diseased and AD donors. Loss of excitatory neurons was greatest in the bands with the highest PHF1 intensity across all neurons (Figure 4C; ρ = -0.82, 95% CI -0.94 to -0.42; permutation *p* = 0.006), and the most affected band (10– 20% depth, corresponding approximately to layer 2) had 40% lower excitatory neuron density in AD than in controls. This relationship was specific to excitatory neurons. Inhibitory neurons showed no detectable loss and no association with PHF1 intensity across the same bands (Figure 4D; ρ = -0.15, 95% CI -0.78 to 0.73; permutation *p* = 0.68). Together, these data indicate that excitatory neuron loss in AD is concentrated in the neuronal populations and cortical depths with the greatest phosphorylated tau burden.

### Nuclear morphological differences in and around tangle-bearing neurons

Tau deforms the nuclear envelope^32^ and relaxes heterochromatin^33^, so we asked whether PHF1_high_ neurons and nearby PHF1_low_ neurons show altered nuclear morphology using the IMC data^28^. PHF1_high_ excitatory neurons had larger, more eccentric nuclei than PHF1_low_ neurons (Figures 5A and S5A; mean area 83.4 versus 65.9 μm^2^) in the absence of a significant difference in total DNA (Figure 5B), consistent with nuclear envelope deformation^32^, swelling or chromatin relaxation^33^. Nuclei were also larger in tangle-bearing versus tangle-free neurons in the CosMx dataset (Figure S5B). PHF1_low_ neurons also had larger nuclei with increasing proximity to the nearest PHF1_high_ neuron, although these remained smaller than those of PHF1_high_ neurons (Figures 5A and S5B). Tangle-bearing neurons also had elevated rRNA stain intensity in the CosMx data (Figure 5C). Tangle-free neurons close to tangles had higher rRNA stain intensity than those further away but did not reach the levels of tangle-bearing neurons (Figure 5C). Representative IMC nuclei from PHF1_high_ and PHF1_low_ neurons, matched for excitatory subtype, illustrate the difference in nuclear size (Figure 5D).

**Figure 5.**
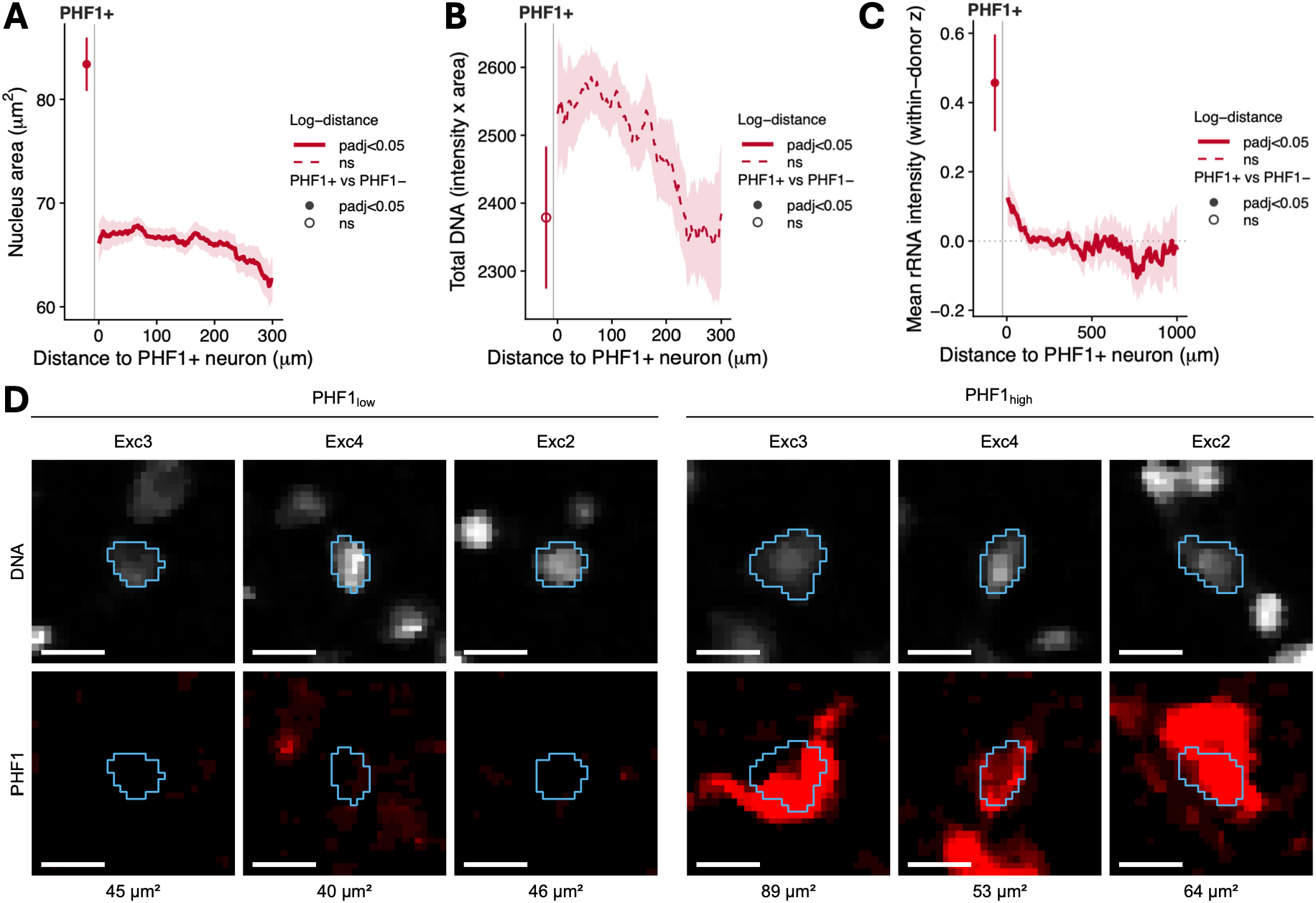
Tangle-bearing neurons have larger nuclei without a detectable difference in total DNA. **(A)** Nucleus area of PHF1_low_ excitatory neurons as a function of distance to the nearest PHF1_high_ neuron (imaging mass cytometry; *n* = 26,653 cells, 44 AD donors). Line and ribbon, 75 μm rolling mean ± 1.96 × s.e.m.; the nucleus area of PHF1_high_ neurons is plotted to the left (mean ± 95% CI of donor means). Nucleus area was greater closer to PHF1_high_ neurons: β = +1.20 μm^2^ per s.d. closer in log-distance (95% CI 0.65, 1.76; adjusted *p* = 4.7 × 10^-5^; linear mixed model, ∼ log-distance + sex + age + PMI + (1|donor), distance capped at 300 μm). PHF1_high_ nuclei are themselves 26.7% larger (17.5 μm^2^ difference [13.2, 21.7], *d_z_* = 1.25, adjusted *p* = 2.3 × 10^-9^; two-sided Wilcoxon signed-rank on donor means, *n* = 44). Solid line, adjusted *p* < 0.05 for log-distance; filled point, adjusted *p* < 0.05 for PHF1_high_ versus PHF1_low_. **(B)** Total nuclear DNA signal (^191^Ir signal intensity × nucleus area) in the same cells and donors, modelled as in **A**. Neither effect is significant: β = -7.29 per s.d. closer (95% CI –32.60, +18.01; adjusted *p* = 0.57) and PHF1_high_ versus PHF1_low_ paired difference -113.7 ([-261.0, +33.6]; adjusted *p* = 0.18). Increased nucleus area in **A** therefore occurs without elevated DNA content. **(C)** rRNA stain intensity in tangle-free Exc-IT-L2-3-*CBLN2*-*HOPX* neurons against distance to the nearest tangle-bearing neuron (CosMx mean rRNA channel, *n* = 24,672 tangle-free neurons, 9 donors). Plotted as uncentred within-donor *z* of log(intensity). rRNA stain intensity was higher towards tangle-bearing neurons: β = +0.039 *z* per s.d. log-distance closer (95% CI 0.025, 0.053; adjusted *p* = 3.6 × 10^-7^, linear mixed model: ∼ log-distance + sex + age + PMI + (1|donor)). Tangle-bearing neurons are themselves elevated by 0.46 ([0.32–0.60]; adjusted *p* = 1.6 × 10^-9^, equivalent to a 1.3-fold raw increase; linear mixed model: ∼ PHF1 status + sex + age + PMI + (1|donor)). *n* = 26,585 neurons, 188 tangle-bearing; 9 donors. Curve and encodings as in **A**. **(D)** Representative IMC crops of PHF1_high_ and PHF1_low_ excitatory neurons, paired by excitatory subcluster (Exc2-4) and ROI. Top, DNA (^191^Ir); bottom, PHF1; outline, segmented nucleus mask for the central nucleus; values, measured nuclear area. Scale bars, 10 μm.

### Glial reactive states vary with distance to tangle-bearing neurons

Having established that proximity to tangle-bearing neurons is associated with transcriptional differences in specific excitatory neuronal populations, we asked whether glial cell states also varied with proximity to tangle-bearing neurons. We first measured the distance from each tangle-bearing neuron to the nearest cell of each glial type and compared it with a null in which tangle-bearing status was randomly reassigned among neurons of the same subtype within each donor (1,000 permutations). Glial distance to tangle-bearing neurons did not differ meaningfully from glial distance to tangle-free neurons (Figure 6A).

**Figure 6.**
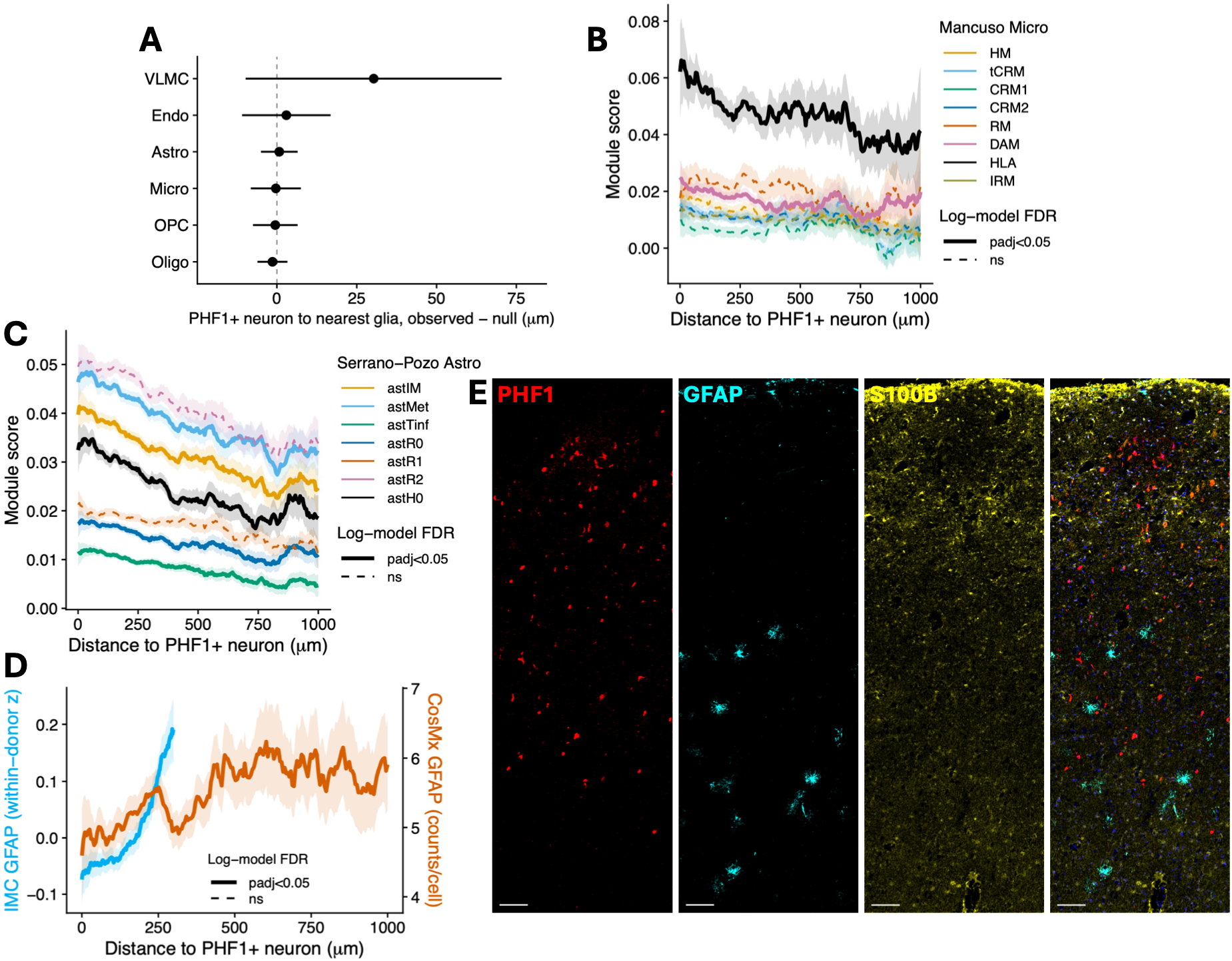
Astrocyte and microglial reactive modules vary with distance to tangle-bearing neurons. **(A)** Distance from each tangle-bearing neuron to the nearest cell of the indicated glial type, observed minus the label-shuffled null. Points, Δ = observed mean minus the median of 1,000 shuffles in which tangle-bearing status is permuted within cell type and donor; lines, 95% confidence intervals calibrated on the permutation null. All adjusted *p* ≥ 0.59, *n* = 9 donors, 398 tangle-bearing neurons. **(B)** Reference microglial signature scores^35^ (Seurat AddModuleScore) in glia within 1,000 μm of a tangle-bearing neuron, modelled per signature against natural-log distance to the nearest tangle-bearing neuron (linear mixed model; adjusted for age, sex, PMI, technical covariates and donor; *n* = 9). The DAM (β = -1.5 × 10^-3^ per s.d. log-distance, 95% CI -2.3 × 10^-3^ to -6.1 × 10^-4^; adjusted *p* = 0.0066) and HLA gene sets (β = -2.8 × 10^-3^, -4.8 × 10^-3^ to -7.1 × 10^-4^; adjusted *p* = 0.034) scored higher in microglia closer to tangle-bearing neurons. Expression of the other microglial modules did not significantly vary with proximity. Lines, observed 75 μm unadjusted rolling mean (shading, 95% CI); colour, module; solid lines, BH-adjusted *p* < 0.05; dashed lines, not significant. *n* = 6,524 microglia; 9 donors. **(C)** Reference astrocyte signature scores^34^ (Seurat AddModuleScore) in glia within 1,000 μm of a tangle-bearing neuron, modelled per signature against natural-log distance to the nearest tangle-bearing neuron (linear mixed model; adjusted for age, sex, PMI, technical covariates and donor; *n* = 9). With increasing proximity to tangle-bearing neurons, astrocytes showed significantly elevated expression of the astH0 (β = -1.6 × 10^-3^ per s.d. log-distance, 95% CI -2.1 × 10^-3^ to -1.1 × 10^-3^; adjusted *p* = 3.2 × 10^-8^), astMet (β = -1.1 × 10^-3^, -1.6 × 10^-3^ to -5.9 × 10^-4^; adjusted *p* = 5.9 × 10^-5^), astIM (β = -1.0 × 10^-3^, -1.5 × 10^-3^ to -5.1 × 10^-4^; adjusted *p* = 1.3 × 10^-4^), astTinf (β = -5.4 × 10^-4^, -8.3 × 10^-4^ to -2.6 × 10^-4^; adjusted *p* = 3.6 × 10^-4^), and astR0 gene modules (β = -4.4 × 10^-4^, -7.6 × 10^-4^ to -1.1 × 10^-4^; adjusted *p* = 0.012). Expression of astR1 and astR2 did not significantly differ with proximity (astR1: β = +1.6 × 10^-4^, -1.7 × 10^-4^ to +4.9 × 10^-4^; adjusted *p* = 0.39; astR2: β = -1.7 × 10^-4^, -6.9 × 10^-4^ to +3.5 × 10^-4^; adjusted *p* = 0.51). Lines, observed 75 μm unadjusted rolling mean (shading, 95% CI); colour, module; solid lines, BH-adjusted *p* < 0.05; dashed lines, not significant. *n* = 16,552 astrocytes; 9 donors. **(D)** GFAP in astrocytes versus distance to the nearest PHF1_high_ or tangle-bearing neuron. Cyan, GFAP protein within 300 μm (IMC; *n* = 26,832 astrocytes, 44 AD donors); orange, *GFAP* RNA within 1,000 μm (CosMx; *n* = 16,552 astrocytes, 9 donors). Lines, 75 μm sliding-window means ± 1.96 s.e.m. Protein: -0.054 *z* per s.d. closer (95% CI -0.067 to -0.042), *p* = 9.4 × 10^-19^; RNA: log_2_FC -0.064 (-0.085 to -0.043) per s.d. closer, adjusted *p* = 1.3 × 10^-6^. GFAP is lower near tangles. **(E)** Representative IMC region of interest spanning the pial surface (top) to the white matter boundary (bottom). Red, PHF1; cyan, GFAP; yellow, S100B; navy, DNA. Scale bar, 100 μm.

We next scored microglia and astrocytes for published gene sets pertaining to cell states^34,35^ (AddModuleScore) and assessed variation in module score with proximity to tangle-bearing neurons. Disease-associated microglia (DAM) and antigen-presenting response (HLA) signatures scored higher in microglia closer to tangle-bearing neurons (Figure 6B). Stably expressed genes showed no detectable distance gradient (Figure S2B; β = -1.7 × 10^-4^, adjusted *p* = 0.52). Astrocytes had higher scores for the astH0, astIM, astR0, astMet and astTinf gene modules with increasing proximity to tangle-bearing neurons, whereas astR1 and astR2 states, defined by canonical *GFAP* upregulation and HSF1-mediated chaperone induction^34^, did not vary with distance (Figure 6C). Stably expressed genes again showed no detectable distance gradient (Figure S2C; β = -5.9 × 10^-5^; adjusted *p* = 0.52; 4–13% of the reported effects). Consistent with the lack of significant astR1 or astR2 gradients, GFAP was reduced in astrocytes proximal to tangles at both the RNA and protein level (Figures 6D and 6E).

Adjustment for cortical depth and proximity to Aβ plaques changed the IMC estimates by only a small amount (Figures S6A–S6C). Of cells within 50 μm of a PHF1_high_ neuron, 19.8–21.9% lay within 40 μm of a 4G8-positive plaque, compared with 14.0% of cells more than 200 μm away (a difference of 5.8–7.9 percentage points; *p* = 0.002–0.006). Adjusting for this small imbalance moved every non-amyloid gradient by less than 0.25 standard errors (for example, AT8 by 0.7% and GFAP by 1.9%). The only large shifts were in 4G8 and H31L21 (20–23%), the channels that themselves report amyloid. Stratifying by plaque proximity gave the same answer: in cells with no plaque within 40 μm, the PHF1_high_ gradients matched the full-dataset estimates (AT8 0.145 vs 0.145; GFAP -0.055 vs -0.054), and no PHF1_high_-distance × plaque-proximity interaction survived FDR correction (0/11; Figures S6D and S6E).

With one exception, scores for the Reactome cell death, stress and autophagy modules examined did not vary with proximity in astrocytes and microglia; only regulated necrosis in astrocytes had a significant log-distance term (β = 7.7 × 10^-4^ per s.d. log-distance, 95% CI 3.4 × 10^-4^ to 1.2 × 10^-3^, adjusted *p* = 0.0026; lower near tangles; Figures S7A–S7F).

## Discussion

The link between neurofibrillary tangle formation and neuronal dysfunction has proved difficult to establish at the single-cell level, and the cells sharing the immediate microenvironment of a tangle-bearing neuron remain largely uncharacterised. Here we combined 6,000-plex spatial transcriptomics with post-run PHF1 immunofluorescence of the same sections and IMC of a larger, separately reported cohort^28^, measuring every cell’s distance to the nearest tangle-bearing neuron and ordering layer 2-3 excitatory neurons along a tangle-associated transcriptional trajectory. Tangle-bearing layer 2-3 excitatory neurons were concentrated towards the end of a continuous trajectory that recapitulated the signature reported in isolated AT8-positive somas of the same laminar identity^26^, marked by higher expression of genes annotated to microtubule regulation, synaptic function and V-ATPase subunits, and elevated p62 and K48-linked ubiquitin at the protein level. Tangle-free neurons of the same subtype varied continuously with distance to the nearest tangle-bearing neuron, largely reproducing the tangle-associated signature not accounted for by their own subthreshold PHF1 immunoreactivity. Local microglia and astrocytes occupied reactive states graded by the same distance, although glial position did not differ detectably.

Elevated p62 alongside K48-linked conjugates, together with increased LAMP1, implicates the endolysosomal-autophagic compartment, which is disrupted early in AD^36,37^ and enriched in tangle-bearing neurons^27^. Impaired clearance and increased pathway induction converge on the same steady-state accumulation and cannot be distinguished in fixed tissue, where flux is not measurable. Interpretation of p62 is further complicated by its transcriptional induction as an NFE2L2 target, a response also elevated in these neurons. The co-occurrence of K48-linked conjugates and phosphorylated tau is nonetheless consistent with a high proteostatic burden, and the presence of these neurons shows that this state is compatible with survival, although whether it would have progressed to loss cannot be determined.

Tangle-free neurons of the same subtype also carried higher subthreshold PHF1 signal at closer proximity, raising the possibility that the gradient reflects early tau accumulation in cells on their way to forming an inclusion. Adjusting for per-cell PHF1 intensity did not attenuate the gradient, indicating that transcriptional state is not accounted for by the tangle-free neuron’s own PHF1 reactivity. Discovery-based differential expression and pre-specified Reactome module scoring converged. Chaperone, stress-response and selective autophagy module scores were higher closer to tangle-bearing neurons, alongside cytoskeletal, synaptic and developmental pathways. Most of these modules were shared with tangle-bearing neurons. However, four did not differ significantly between tangle-bearing and tangle-free neurons: hypoxia response, reactive oxygen species detoxification, cytoprotection by *HMOX1* and mitophagy were graded in proximal tangle-free neurons but did not differ significantly in tangle-bearing neurons. Chaperone engagement in tauopathy has been studied almost exclusively in the aggregate-bearing cell, where HSP90 and HSF1 activity has been argued both to triage tau and to permit its accumulation^38^. The proximity-associated elevation of these scores here in neurons with no detectable inclusion shows that chaperone-related transcriptional pathways are not confined to the aggregate-bearing cell. The same holds for aggrephagy module scores; elevated selective autophagy transcripts in a neuron without an inclusion are consistent with substrate below the detection threshold of PHF1 immunofluorescence or a signal originating outside the cell.

Proximity to tangle-bearing neurons was associated with glial state but not detectably with glial location. Microglia had higher DAM and HLA^35^ scores with proximity, and astrocytes had higher homeostatic, intermediate and early reactive module scores rather than the classical reactive profile. GFAP protein and RNA were lower near tangle-bearing neurons and the *GFAP*-high astR1 and astR2 states^34^ were not associated with proximity to tangles, although reactive astrocytes do not uniformly have elevated GFAP^39^. Reactive astrocyte markers are higher near plaques^34^, suggesting that tangle and plaque niches are associated with distinct astrocyte states. The terminal astMet and astTinf states, which in the reference dataset vary with disease stage and in the case of astMet have been interpreted as exhausted^34^, scored higher near tangle-bearing neurons. Distance to a tangle may therefore capture, at cellular scale, the cumulative exposure to pathology that has been inferred across Braak stages.

Our study has several limitations. Because each section captures a single time point, a proximity gradient cannot establish the direction of the relationship, as tangle-free neurons may be responding to the tangle-bearing neuron, or both may reflect a local factor that preceded either. Vascular architecture and amyloid burden were not modelled in the spatial transcriptomics analyses, and we were only able to observe neurofibrillary tangle proximity in two dimensions. The transcriptomic cohort included tissue from only nine donors, which limited our power to test whether these gradients differ by disease stage. Tangle status was assigned by manual annotation of somatic PHF1 immunoreactivity, which identifies inclusions but does not resolve soluble or oligomeric tau; neurons scored as tangle-free may carry species below this threshold, and the gradient in subthreshold PHF1 intensity we report indicates that some do. This design is agnostic to which tau species the gradient is structured around. Repeating it with a conformation-specific reporter of soluble or oligomeric tau, in place of or alongside a phospho-tau antibody such as PHF1, would establish whether transcriptional state is structured around the fibrillar inclusion or around the soluble pool that surrounds it, and would distinguish a tangle-centred gradient from one centred on a species the tangle merely marks. Finally, transcript abundance is an imperfect proxy for protein activity, particularly for the 14-3-3 proteins, whose function is set by target phosphorylation rather than by expression level.

One hypothesis these findings raise is that tangle-bearing neurons, or the cellular state they mark, are a source of injury to the tissue around them. The population in which we resolve this gradient, Exc-IT-L2-3-*CBLN2*-*HOPX*, includes calbindin- and reelin-expressing excitatory neurons previously identified as selectively vulnerable in AD^28,29^, placing the gradient in cells already implicated in early entorhinal degeneration. In the IMC cohort, excitatory neuron loss was greatest in the clusters lying closest to PHF1_high_ neurons and at the cortical depths with the most phosphorylated tau, whereas inhibitory neurons were not detectably depleted. Because tangle-free neurons within range of a tangle far outnumber tangle-bearing neurons, this would reconcile the excess of neuronal loss over tangle number^19,20^ and the persistence of tangle-bearing neurons^21–24^ with the close relationship between tangle burden and clinical severity^18^.

A cross-sectional study cannot establish causality, but model systems can. Whether the tangle-associated gradient requires a persisting source is tractable in tauopathy models, where tau expression in inclusion-bearing neurons can be suppressed and the neighbouring state measured before and after. Whether it contributes to neuronal loss can be tested by following the survival of neurons surrounding tangles as a function of their distance from them, as has been done for tangle-bearing neurons themselves^21–23^. The answer to this second question has the clearest translational bearing. If the neighbour state contributes to loss, the therapeutic implication is combinatorial, since lowering tau and protecting surviving neighbours would address different arms of the same problem. If it instead marks neurons on a route to loss, it is a marker of risk rather than a target, and tau-directed therapies would need to be evaluated by the state of neurons around inclusions as well as by the inclusion burden.

The cellular differences associated with tangles extend beyond the neurons that bear them, to nearby tangle-free neurons and local glia graded by distance, and into the neuronal populations and cortical depths in which loss in AD is concentrated. Selective neuronal vulnerability may therefore reflect not only a neuron’s susceptibility to forming a tangle, but also where it lies relative to one.

## Resource availability

### Lead contacts

Professor Paul Matthews

Dr Anna Mallach

### Materials availability

This study did not generate unique reagents.

### Data and code availability

Raw CosMx data have been deposited on GEO (GSE349866), and processed CosMx data, source data and statistics logs at Zenodo (https://doi.org/10.5281/zenodo.23188273); both are publicly available as of the date of publication. Raw and processed IMC data were previously deposited at figshare (see Key Resources). All original code has been deposited at Zenodo (https://doi.org/10.5281/zenodo.23188012). Any additional information required to reanalyse the data reported in this paper is available from the corresponding authors upon request.

## Acknowledgements

We thank the study participants, their families, and the coordinating brain banks, without whom this study would not have been possible. We thank all members of the Matthews laboratory at the UK Dementia Research Institute at Imperial College London for discussions and feedback.

SLB is supported by the Alzheimer’s Research UK Patricia Wood-Smith PhD Scholarship (ARUK-PhD2022-026). AC was supported by an Alzheimer’s Society career development grant (628; AS-CD-23-004) and the European Union’s Horizon 2020 research and innovation programme under the Marie Skłodowska-Curie award (101146149). NNF and AM were both supported by the Edmond J Safra Foundation as part of their Edmond and Lily Safra Fellowships. PMM gratefully acknowledges support from the UK Dementia Research Institute and personal funding of his chair by the Edmond J Safra Foundation and Lily Safra.

## Author contributions

Conceptualisation - SLB, AM, PMM; Formal analysis - SLB; Funding acquisition - PMM; Investigation - SLB, NL, AC; Methodology - SLB, NL, MT, NNF; Supervision - JSJ, EPD, AM, PMM; Validation - SLB, JW, MP, SMP, EA; Visualisation - SLB; Writing (original draft) - SLB, AM, PMM; Writing (review and editing) - SLB, NL, MP, EA, MT, JSJ, EPD, AM, PMM. All authors approved the final manuscript.

## Declaration of interests

PMM received research funding from Biogen and Nimbus Therapeutics and has acted as a consultant to GlaxoSmithKline, Biogen, Astex/Otsuka, Nimbus and Sudo Therapeutics. He is jointly employed by Imperial College London and the Rosalind Franklin Institute. The remaining authors declare no competing interests.

## Supplemental information titles

**Figure S1.** The distance gradient in tangle-free Exc-IT-L2-3-CBLN2-HOPX neurons is not explained by PHF1 intensity.

**Figure S2.** Stably expressed genes show no comparable spatial gradient.

**Figure S3.** Robustness of the tangle-proximity transcriptional gradients.

**Figure S4.** Apparent cell death module enrichment in and around tangle-bearing neurons is driven by genes encoding the anti-apoptotic 14-3-3 scaffold, not by apoptotic effectors.

**Figure S5.** Nucleus eccentricity and CosMx replication of nuclear enlargement.

**Figure S6.** Adjustment for Aβ plaque proximity and cortical depth.

**Figure S7.** Cell death, stress and autophagy modules in glia did not vary detectably with tangle proximity.

**Table S1.** CosMx and IMC donor characteristics

**Table S2.** Cell annotation marker gene sets

**Table S3.** Cell quality control by donor

**Table S4.** Genes associated with pseudotime along the tangle-associated continuum in Exc-IT-L2-3-CBLN2-HOPX neurons

**Table S5.** GO enrichment of the Table S4 genes (padj < 0.1 and |logFC_trajectory| > 0.25)

**Table S6.** Gene expression vs distance to the nearest tangle-bearing neuron, tangle-free Exc-IT-L2-3-CBLN2-HOPX neurons

**Table S7.** GO enrichment of the Table S6 genes (padj < 0.1)

**Table S8.** Exclusion-radius sensitivity

## STAR Methods

### Key resources table

| REAGENT or RESOURCE | SOURCE | IDENTIFIER |
|---|---|---|
| Antibodies |  |  |
| PHF1 anti-tau monoclonal antibody | Peter Davies Laboratory, Albert Einstein College of Medicine | RRID:AB_2315150 |
| Alexa Fluor 568 donkey anti-mouse IgG (H+L) | Thermo Fisher Scientific | Cat# A10037; RRID:AB_11180865 |
| Chemicals, peptides, and recombinant proteins |  |  |
| 10% NBF | Molecular Dimensions | BS-0900E-2500ML |
| PBS | Thermo Fisher Scientific | AM9625 |
| SDS | Thermo Fisher Scientific | AM9822 |
| Ethanol | Thermo Fisher Scientific | 16695992 |
| Tris | Sigma-Aldrich | 10708976001 |
| Glycine | Sigma-Aldrich | G7126 |
| DEPC-treated water | Thermo Fisher Scientific | AM9922 |
| 20X SSC (DNase, RNase free) | Thermo Fisher Scientific | AM9763 |
| Sulfo-NHS-Acetate powder | Thermo Fisher Scientific | 26777 |
| Xylene | Merck Life Science UK Limited | 534056-4L |
| 100% Deionized formamide | Thermo Fisher Scientific | AM9342 |
| Donkey serum | Abcam | Cat# ab7475; RRID:AB_2885042 |
| TrueBlack<sup>®</sup> Autofluorescence Quencher | Cell Signaling Technology | 92401S |
| DAPI | Thermo Fisher Scientific | 10116287 |
| ProLong<sup>TM</sup> Diamond Antifade Mountant | Invitrogen | P36970 |
| Critical commercial assays |  |  |
| CosMx Human 6K Discovery Panel | Bruker Spatial Biology | 531-121500041 |
| CosMx RNA 6Kplx 2sl Img Tray Kit | Bruker Spatial Biology | 531-122000165 |
| CosMx Human Neuroscience Cell Segmentation Kit, Ch1-4 (RNA) | Bruker Spatial Biology | 531-121500043 |
| CosMx RNAse Inhibitor (2-Slide Run) | Bruker Spatial Biology | 531-121500004 |
| CosMx Cleaning Tray | Bruker Spatial Biology | 531-122000152 |
| CosMx FF Slide Prep Kit (RNA) | Bruker Spatial Biology | 531-121500007 |
| CosMx RNA Instrument Buffer Kit (Medium) | Bruker Spatial Biology | 531-100481 |
| CosMx Flow Cell Assembly Kit | Bruker Spatial Biology | 531-122000061 |
| Deposited data |  |  |
| Raw CosMx data | Gene Expression Omnibus | GSE349866 |
| Processed CosMx data | Zenodo | https://doi.org/10.5281/zenodo.23188273 |
| snRNA-seq reference data<sup>28</sup> | Gene Expression Omnibus | GSE333997 |
| Raw IMC data<sup>28</sup> | figshare | https://doi.org/10.6084/m9.figshare.32532627 |
| Processed IMC data<sup>28</sup> | figshare | https://doi.org/10.6084/m9.figshare.32526492 |
| Software and algorithms |  |  |
| Original analysis code | This paper | https://doi.org/10.5281/zenodo.23188012 |
| R v4.4.3 (CosMx) and v4.4.2 (IMC) | R Core Team | RRID:SCR_001905 |
| Seurat v5.3.1 (SeuratObject v5.4.0, sctransform v0.4.2) | Hao et al., 2024<sup>40</sup> | RRID:SCR_016341 |
| SingleCellExperiment v1.28.1 and SummarizedExperiment v1.36.0 | Amezquita et al., 2020<sup>41</sup> | RRID:SCR_026794 |
| limma v3.62.2 (with statmod v1.5.2) | Ritchie et al., 2015<sup>42</sup> | RRID:SCR_010943 |
| edgeR v4.4.2 | Chen et al., 2025<sup>43</sup> | RRID:SCR_012802 |
| variancePartition v1.36.2 | Hoffman and Roussos, 2021<sup>44</sup> | RRID:SCR_019204 |
| lme4 v1.1-38 | Bates et al., 2015<sup>45</sup> | RRID:SCR_015654 |
| lmerTest v3.2-0 | Kuznetsova et al., 2017<sup>46</sup> | RRID:SCR_015656 |
| enrichR v3.4 | Kuleshov et al., 2016<sup>47</sup> | RRID:SCR_001575 |
| AUCell v1.28.0 | Aibar et al., 2017<sup>48</sup> | RRID:SCR_021327 |
| ComplexHeatmap v2.22.0 | Gu et al., 2016<sup>49</sup> | RRID:SCR_017270 |
| ggplot2 v4.0.3 | Wickham, 2016 | RRID:SCR_014601 |
| dplyr v1.2.1 | Wickham et al. | RRID:SCR_016708 |
| qs v0.27.3 | Ching | https://doi.org/10.32614/CRAN.package.qs |
| Matrix v1.7-6 | Bates et al. | https://doi.org/10.32614/CRAN.package.Matrix |
| argparse v2.3.1 | Davis | https://doi.org/10.32614/CRAN.package.argparse |
| zoo v1.9-0 | Zeileis and Grothendieck, 2005<sup>50</sup> | https://doi.org/10.32614/CRAN.package.zoo |
| glmGamPoi v1.18.0 | Ahlmann-Eltze and Huber, 2021<sup>51</sup> | https://doi.org/10.18129/B9.bioc.glmGamPoi |
| scDblFinder v1.23.4 | Germain et al., 2021<sup>52</sup> | RRID:SCR_022700 |
| scMerge v1.22.0 | Lin et al., 2019<sup>53</sup> | https://doi.org/10.18129/B9.bioc.scMerge |
| RANN v2.6.2 | Jefferis et al. | https://doi.org/10.32614/CRAN.package.RANN |
| uwot v0.2.4 (UMAP) | McInnes et al., 2018<sup>54</sup> | https://doi.org/10.32614/CRAN.package.uwot |
| Harmony v1.2.4 | Korsunsky et al., 2019<sup>55</sup> | RRID:SCR_022206 |
| Slingshot v2.14.0 | Street et al., 2018<sup>56</sup> | RRID:SCR_017012 |
| Steinbock v0.16.3 | Windhager et al., 2023<sup>57</sup> | https://github.com/BodenmillerGroup/steinbock |
| Python v3.11 | Python Software Foundation | RRID:SCR_008394 |
| NumPy v2.2.0 | Harris et al., 2020<sup>58</sup> | RRID:SCR_008633 |
| tifffile v2025.10.16 | Gohlke | https://doi.org/10.5281/zenodo.6795860 |
| scikit-image v0.25.2 | van der Walt et al., 2014<sup>59</sup> | RRID:SCR_021142 |
| AtoMx Spatial Informatics Platform v1.3.2 | Bruker Spatial Biology | https://www.bruker.com |
| napari v0.4.19.post1 with napari-cosmx v0.4.17.3 and affinder v0.5.0 | Sofroniew et al.; Bruker Spatial Biology; Nunez-Iglesias et al. | RRID:SCR_022765 |
| Leica Application Suite X v4.7.0.28176 | Leica Microsystems | RRID:SCR_013673 |
| Other |  |  |
| BOND Plus slides | Leica Biosystems | S21.2113.A |
| CosMx Spatial Molecular Imager | Bruker Spatial Biology | RRID:SCR_023909 |
| STELLARIS 8 (DMI8-CS) | Leica Microsystems | RRID:SCR_024664 |
| Reactome pathway gene sets | Milacic et al. (2024)<sup>60</sup> | RRID:SCR_003485 |
| AT8 gene sets | Otero-Garcia et al. (2022)<sup>26</sup> | 10.1016/j.neuron.2022.06.021 |
| Astrocyte gene sets | Serrano-Pozo et al. (2024)<sup>34</sup> | 10.1038/s41593-024-01791-4 |
| Microglia gene sets | Mancuso et al. (2024)<sup>35</sup> | 10.1038/s41593-024-01600-y |

### Experimental model and study participant details

#### Ethical approval

This study was carried out in accordance with the Regional Ethics Committee and Imperial College London Use of Human Tissue guidelines. All the brain banks used have generic research ethics committee approval to function as research tissue banks and therefore did not require additional ethics panel approvals for use by UK researchers.

### CosMx tissue cohort

Frozen EC samples (*n* = 9) for spatial transcriptomics were obtained from three UK brain banks: Oxford Brain Bank (University of Oxford; *n* = 2), South West Dementia Brain Bank (University of Bristol; *n* = 4), and the London Neurodegenerative Diseases Brain Bank (King’s College London; *n* = 3). Samples were selected based first on neuropathological diagnosis of AD. Three Braak stage I samples without a diagnosis of AD were selected based on PHF1 positivity in the previously reported cohort^28^. The final cohort comprised three donors per pathology group: Braak stage I without AD but positive for PHF1 (*n* = 3), Braak stage IV with AD (*n* = 3) and Braak stage VI with AD (*n* = 3). Five donors were female and four male; median age at death was 85 years (range 69–98) and median *post-mortem* interval was 26.8 hours (range 3.0–85.8). Donor characteristics are given in Table S1. We excluded cases with clinical or *post-mortem* diagnosis of potentially confounding neurological conditions, including but not limited to Parkinson’s disease, Huntington’s disease, progressive supranuclear palsy, epilepsy, hippocampal sclerosis, frontotemporal dementia, age-related tau astrogliopathy, argyrophilic grain disease, and traumatic brain injury. Further exclusion criteria included type 2 diabetes mellitus, the presence of infarcts greater than 10 mm in diameter unless confined to the cerebellum, and cerebrovascular disease (such as cerebral atherosclerosis, small vessel disease, arteriosclerosis) described as moderately severe to severe. A tenth section (Braak stage V, AD) was imaged but excluded from all analyses after cell type annotation because the tissue was subsequently found to consist of white matter rather than EC grey matter.

### IMC tissue cohort

The previously reported IMC cohort^28^ consisted of *post-mortem* EC sections of 8 μm thickness from 18 non diseased controls (Braak 0–I; 11 female, 7 male; median age 72.5 years, range 67– 90; 54 ROIs) and 44 donors with AD (Braak stage III–IV, *n* = 19; Braak VI, *n* = 25; 23 female, 21 male; median age 84 years, range 68–101) from five brain banks (Cambridge Brain Bank, South West Dementia Brain Bank, Oxford Brain Bank, Multiple Sclerosis and Parkinson’s Tissue Bank and the Netherlands Brain Bank; Table S1). The cohort had a median *post-mortem* interval of 21.1 hours (range 3.0–41.0). Three regions of interest were ablated per donor, giving 186 ROIs and 170,195 annotated single cells across 35 channels for the full 62 donor cohort, and 132 ROIs and 127,157 cells for the AD donors only. Donors were drawn from a larger IMC series (67 donors, 195 ROIs) described previously^28^; donors with fewer than three ROIs were excluded. Tissue sourcing, antibody panel design, staining and acquisition were as reported there. The work presented here used the existing cell type and tangle annotations. PHF1_high_ annotations in the published object were obtained via Otsu thresholding of PHF1 intensity. Unless otherwise stated, IMC analyses were restricted to the 44 donors with AD; the non-diseased donors were used only as the reference for neuronal loss.

## Method details

### CosMx spatial molecular imaging

Fresh-frozen EC sections were cut at 10 μm, mounted onto Leica BOND Plus slides and stored at -80 °C until processing. On day one, sections were fixed in pre-cooled 10% neutral buffered formalin (NBF) for 2 hours at 4 °C. Following fixation, slides were washed in 1× phosphate buffered saline (PBS, pH 7.4) and baked for 30 minutes at 60 °C. Sections were then washed three times in 1× PBS, once in 4% sodium dodecyl sulphate (SDS), and a further three times in 1× PBS. Slides were subsequently rehydrated through a graded ethanol series (50%, 70%, and 100%) and air dried for 10 minutes at room temperature.

Heat-induced target retrieval was performed at 100 °C in 1× target retrieval solution for 15 minutes. Slides were washed in DEPC-treated water followed by 100% ethanol and air dried for 30 minutes at room temperature. An incubation frame was applied to each slide prior to tissue permeabilisation. RNA targets were exposed by incubation with Proteinase K at 3 µg/mL at 40 °C for 30 minutes, followed by washes in 1× PBS.

Fiducial markers were applied at a final concentration of 0.00015% for 5 minutes at room temperature to provide stable reference points for accurate spatial image registration and alignment across multiple rounds of fluorescent probing and imaging. Sections were washed in 1× PBS, fixed in 10% NBF, and washed twice in NBF stop buffer (0.1 M Tris, 1.1 M glycine in DEPC-treated water). To prevent non-specific binding, 100 mM sulfo-NHS-acetate was applied for 15 minutes at room temperature, followed by a series of washes in 2× saline-sodium citrate (SSC) buffer.

Slides were incubated overnight (16 to 18 hours) at 37 °C in a hybridisation oven with the Bruker Human 6K Discovery Panel, comprising approximately 6,000 target-specific *in situ* hybridisation probes coupled to unique photocleavable oligonucleotide readout barcodes, together with an rRNA segmentation marker^61^. Following in situ hybridisation, sections underwent two stringent washes in 50% 4× SSC/deionised formamide at 37 °C for 25 minutes, followed by 2× SSC, to remove off-target probes.

Nuclei were stained with DAPI for 15 minutes. A fluorescently labelled histone marker was then applied for cell segmentation and incubated for 1 hour at room temperature. The flow cell was assembled using a Bruker flow cell coverslip, and 2× SSC buffer was used to prevent the tissue from drying out during imaging.

Prepared flow cells were loaded onto the CosMx Spatial Molecular Imager using Bleaching Protocol B (90 seconds) and Cell Segmentation Profile B. A total of 605 fields of view (FOVs) across 2 slides were selected for imaging.

### Post-CosMx immunofluorescence staining

Slides were soaked in xylene overnight, before the flow cell cover was carefully removed with a blade and the coverslip taped to avoid glass falling onto the slide. Slides were submerged in 2× saline-sodium citrate (SSC) three times for five minutes each, then blocked for 30 minutes with 10% donkey serum in phosphate buffered saline (PBS) with 0.3% Triton X-100. 200 *μl* of PHF1 antibody diluted at 1:5000 was applied and left on the slides overnight at 4 °C in a dark humidified chamber. The following morning, the slides were washed in PBS three times for five minutes each, then the Alexa Fluor 568 donkey anti-mouse secondary antibody, diluted at 1:1000 in PBS, was applied and left in the dark for one hour at room temperature. Slides were then washed in PBS three times for five minutes each, then underwent autofluorescence quenching in 1× TrueBlack for 1 minute. Slides were washed in PBS three times for five minutes each, then incubated in 1:1000 DAPI for 10 minutes. Slides were washed again in PBS for five minutes and then mounted with ProLong Diamond Antifade Mountant and a glass coverslip.

### Post-run immunofluorescence imaging

Slides were imaged using a Leica Stellaris 8 confocal microscope (DMI8-CS stand, LAS X 4.7.0.28176) with an HC PL APO CS2 20×/0.75 DRY objective, pinhole 1 AU, optical zoom 2. DAPI was collected on a HyD S detector and the PHF1 Alexa 568 signal on a HyD X detector. Whole section images were acquired as tile scans and merged in LAS X, giving a 2-channel mosaic per section with an isotropic pixel size of 0.2841 μm.

### Post-run immunofluorescence alignment

Post-run PHF1/DAPI images were registered to the CosMx morphology mosaics in napari. CosMx napari datasets were built with the napari-cosmx plugin (v0.4.17.3; stitch-images), which places all CosMx layers in a common millimetre world frame at 0.1203 *µm* px^-1^. For each section, corresponding nuclei were identified manually in the post-run DAPI and CosMx DAPI layers and a 6-degree-of-freedom affine transform was estimated from these landmark pairs with the affinder plugin (skimage.transform.AffineTransform). Pairs were added iteratively, with the overlay updating live, until nuclear outlines were judged coincident across the whole section by visual inspection. The fitted transforms were near-rigid: a rotation of 90.05–90.36°, a fitted pixel scale of 0.2846 × 0.2851 μm px^-1^, residual anisotropy between the two fitted axis scales ≤ 0.35% and shear ≤ 0.39°, indicating that the landmark fit introduced no appreciable geometric distortion. The exported matrices were inverted and composed analytically with the plugin’s FOV-stitching geometry to map each CosMx cell-segmentation mask into post-run stain pixel coordinates, and PHF1 intensity was aggregated per cell mask without resampling either image.

### Tangle annotation

Tangle-bearing neurons were annotated manually on aligned post-run PHF1/DAPI images. A cell was annotated as tangle-bearing when compact or pyramid-shaped PHF1 immunoreactivity filled the soma, with the inclusion lying within the segmented cell mask. Diffuse or granular PHF1 signal without fibrillar morphology (pre-tangle staining), neuropil threads and dystrophic neurites were not annotated as tangle-bearing. Annotation was performed by one rater blinded to cell type label, donor identity and transcriptomic data. 95^th^ percentile PHF1 intensity was extracted per cell and converted to robust z score within sample. The 100 neurons per donor with the highest PHF1 intensity that had been annotated as tangle-free were then re-evaluated by three raters including the original rater, and 8 additional cells were reclassified as tangle-bearing following agreement between the raters.

### Reliability of tangle annotations

Reliability of the manual tangle annotations was assessed by blinded re-annotation by the original annotator, 33 weeks after the original calls. The 600 neurons comprised 150 tangle-bearing neurons (donor-stratified random draw) and 450 tangle-free neurons (three per tangle-bearing neuron, drawn at random within the same donor and cell type). Each was presented as a 120 μm PHF1/DAPI crop centred on the target, with a fixed intensity scale per section. Cells were identified only by random codes and shown in random order, with no label, donor or cell-type information. Agreement with the analysis labels was summarised as observed agreement, Cohen’s κ^30,62^ and Gwet’s AC1^63^, with sensitivity and specificity against the original calls. 95% CIs were obtained from a donor-level bootstrap (B = 5,000).

Aggregated intensities were concordant with the manual PHF1 scoring performed in napari on the same overlay (ROC AUC 0.93–1.00 per section; median 95^th^-percentile masked intensity 78–216 in tangle-bearing versus 28–51 in tangle-free cells). Intensity could not be extracted for 336 cells (0.2%; none tangle-bearing) whose segmentation mask lay less than half inside the registered PHF1 image at section edges.

### Image display and representative panels

Display panels were rendered from the registered images with the same transform used for quantification, without resampling, and CosMx segmentation polygons were drawn as outlines coloured by cell type. Channels were additively blended (DAPI, blue; PHF1, red) after per-channel background subtraction and linear scaling of each channel to its own 99.5^th^ percentile, with no gamma; scaling is per panel and for display only, and all reported intensities came from the raw images. Whole-section overviews were downsampled 6-fold; single-FOV views and 60 μm single-cell crops are at native resolution. Scale bars were derived from each section’s fitted pixel size; panels were exported at 300 dpi.

## Quantification and statistical analysis

### Cell segmentation and quality control

Cells were segmented in AtoMx using the default method. Quality control was performed based on Bruker guidelines. For each cell, negative-probe and false-code contamination were expressed as a percentage of the gene-panel transcript count. Cells were retained if they carried at least 50 transcripts, if the transcript and detected-gene counts both fell between the 1^st^ and 99.5^th^ percentiles of the pooled distribution, and if negative-probe and false-code percentages were each at or below 2%. Fields of view with a mean of fewer than 100 transcripts per cell were excluded in full. Doublets were then identified with scDblFinder run within donor at an expected doublet rate of 0.03, and predicted doublets removed. 28,288 cells of an initial 207,582 were excluded during quality control, and a further 12,118 were identified as doublets and removed. 167,176 cells were taken forward to analysis.

### Normalisation

Counts were variance-stabilised with SCTransform using the glmGamPoi method, regressing out total transcript count, negative-probe percentage and false-code percentage. Genes detected in fewer than 0.5% of retained cells would have been discarded; however, all 6,175 panel targets passed this threshold and were carried forward. SCT-corrected values were used for cell type annotation, module scoring and marker detection.

### Cell type annotation

Cell identities were assigned by two sequential rounds of marker-based annotation using an EC snRNA-seq reference generated previously by our group^28^. In the first round, reference labels were collapsed to broad classes (astrocytes, microglia, oligodendrocytes, OPCs, endothelial cells, VLMCs, glutamatergic and GABAergic neurons). Reference and query were restricted to their shared genes, and marker sets were defined per class by one-versus-rest Wilcoxon tests on the reference SCT assay (positive markers only, detected in at least 20% of cells, log_2_FC at least 0.25, adjusted *p* < 0.05), taking the 100 genes with the largest fold change per class and requiring at least 10 genes per set. Each CosMx cell was then scored against every marker set with AUCell, using rankings built on SCT-corrected counts. Per-class assignment thresholds were derived from the CosMx AUC distributions (AUCell_exploreThresholds, falling back to the 90^th^ percentile where no threshold was returned) and bounded below at 0.035. Thresholds were then adjusted by the class proportions of the reference, lowering the threshold for common classes and raising it for rare ones. Cells were assigned to the class with the highest ratio of AUC to adjusted threshold, provided that the ratio was at least 1 and the AUC exceeded 0.035; cells meeting no threshold were labelled unassigned. Cells assigned to either neuronal class were carried into a second round against neuronal subtype marker sets from the same reference, collapsed to five excitatory and four inhibitory subtypes. This round used the same procedure with stricter marker selection (log_2_FC ≥ 1, ≤ 60 genes per set) and a lower threshold floor and AUC minimum (0.027). Neurons that did not score highly for any neuronal subtype marker gene set but were annotated as neurons in the first round were annotated as unassigned neurons. Subtype labels replaced the broad neuronal label where assigned. Excitatory subtypes are named by CosMx marker genes distinguishing them and by canonical layer and projection class derived from the corresponding reference population. Of 167,176 cells, 126,414 (75.6%) were assigned a cell type label. Of the 62,742 neurons annotated, 47,209 (75.2%) were assigned a subtype label.

### Distance calculation

For every tangle-free cell, the distance to the nearest tangle-bearing neuron was computed from slide-global segmentation centroids by k-nearest-neighbour search on a k-d tree (RANN), taking the single nearest anchor and converting from millimetres to micrometres. Anchors were defined as tangle-bearing cells assigned to any of the ten neuronal labels, comprising nine confidently assigned neuronal subclusters and one population that was confidently annotated as neuronal in the first labelling round but did not score highly for any individual neuronal subcluster gene set in the second round. The query set comprised all tangle-free cells of any class. Searches were performed independently within each tissue section. Tangle-bearing neurons carry no distance of their own; they are therefore absent from every distance-based model.

### Permutation of tangle-bearing labels

Tangle annotations were shuffled 1,000 times within each cell type and section stratum, holding each stratum’s tangle-bearing count fixed, with distances to the nearest tangle-bearing neuron recomputed per labelling.

### Glial proximity to tangle-bearing neurons

For each glial type, proximity was summarised as the mean distance from a tangle-bearing neuron to the nearest cell of that type, averaged within donor and then over the nine donors, and recomputed identically over the 1,000 permuted tangle annotations. Because labels were shuffled within cell type and donor, laminar position and per-donor tangle counts are matched. The reported statistic is Δ, the observed mean minus the median of the 1,000 permuted values, in μm. Two-sided empirical *p* values were taken against the null distribution and BH-adjusted across the six glial types.

### Construction of the tangle-associated trajectory

Exc-IT-L2-3-*CBLN2*-*HOPX* neurons were embedded on their own with SCTransform (glmGamPoi; regressing library size and negative-probe and false-code fractions), principal component analysis, Harmony integration by donor (first 10 components; elbow of the variance spectrum) and UMAP. Neurons were clustered by *k*-means and pseudotime was inferred with Slingshot, rooted at the cluster (≥100 neurons) with the lowest fraction of tangle-bearing neurons, as the curve-weighted average across lineages. Settings were chosen to maximise the mean per-donor AUC of tangle-bearing versus tangle-free neurons along pseudotime, in donors with ≥10 tangle-bearing neurons (UMAP, *k* = 12; *n* = 6). This AUC was tested against 1,000 within-donor permutations of tangle labels, with the full selection repeated on each permutation. The percentage of tangle-bearing neurons across ten equal-width pseudotime bins was compared by Friedman test with donor as the block.

Pseudotime was modelled against scaled log-distance as for differential expression. The breakpoint at which pseudotime departs from its distal baseline was estimated with a single-hinge model on raw distance (grid search, 10–900 μm), with a 95% CI from a donor bootstrap (999 replicates).

### Differential expression along the tangle-associated trajectory

Genes associated with pseudotime were identified with a cell-level linear mixed model in dream (variancePartition). Genes detected in ≥5% of neurons were tested (*n* = 2,437). Counts were normalised to counts per million and transformed with voomWithDreamWeights. All neurons were modelled as: expression ∼ pseudotime + log_2_ library size + negative-probe fraction + sex + age + *post-mortem* interval + (1|donor). Age and *post-mortem* interval were scaled. Pseudotime entered divided by its s.d. Empirical Bayes moderation was applied and *p* values were adjusted by the Benjamini-Hochberg method. Effect sizes are reported as the log_2_ fold change across the whole trajectory (the per-unit slope × the pseudotime range). Genes were called at adjusted *p* < 0.1 and |log_2_FC| > 0.25. These associations describe genes that are differentially expressed along the tangle-associated axis.

### Distance-dependent differential expression

To test whether transcriptional state varies with proximity to a tangle, tangle-free cells lying within 1,000 μm of the nearest tangle-bearing neuron were modelled against continuous distance. Distance was natural-log transformed and divided by its standard deviation, computed within each cell type after the cap and without centring, so coefficients are expressed per standard deviation of log-distance. Tangle-bearing neurons carry no distance of their own and are absent from these models. Genes detected in at least 5% of the modelled cells of a given cell type were tested. Counts were normalised to counts per million and observation-level precision weights estimated with voomWithDreamWeights. Each gene was then fitted by variancePartition dream as expression ∼ scaled log-distance + log_2_ total counts + negative-probe percentage + sex + age + *post-mortem* interval + (1|donor), with empirical Bayes moderation.

### Robustness of the distance-dependent differential expression analysis

As a first sensitivity analysis, the model was refitted after excluding tangle-free cells lying within 10, 20, 30 and 50 μm of the nearest tangle-bearing neuron. The gene set tested was the same as in the primary analysis.

A second sensitivity analysis, investigating the laminar position of tangle-free neurons as a confound in the contrast, was performed by adding scaled cortical depth into the model as a fixed effect.

Third, tangles lying outside the imaged plane were simulated. Tangles detected within the section were treated as complete. Additional tangles were drawn as a Poisson process at the volumetric density implied by each donor’s observed in-plane tangle density and the depth of tissue the section samples (section thickness plus inclusion diameter, 18 μm). Tangles were simulated over the imaged area and extended to ±500 μm above and below the section, excluding the sampled slice itself so that no simulated tangle could occupy the plane already observed. Simulated tangles were placed by displacing a randomly chosen observed tangle by a Gaussian offset (σ = 200 μm) and retaining those falling within the imaged area, bracketing the extent to which tangles cluster in three dimensions. Each tangle-free cell’s distance was recomputed as the shortest three-dimensional distance to any tangle, observed or simulated. As placement is stochastic, simulation and fitting were repeated ten times per condition and the gene-level coefficients combined across replicates with their between-replicate variance. The gene set tested was the same as in the primary analysis.

### Pathway overrepresentation

Genes significant at BH-adjusted *p* < 0.1 in each differential expression analysis were split by the sign of their coefficient and each direction tested separately for overrepresentation with enrichR against the GO Biological Process, Cellular Component and Molecular Function 2025 libraries. For the tangle-associated trajectory analysis, gene expression also had to change by more than 0.25 log_2_ units across the whole trajectory. The background was the set of genes tested in that analysis. Terms were retained if they contained between 15 and 250 genes in the GO library, overlapped the query list by at least three genes (five for the tangle-associated trajectory analysis), and returned an odds ratio above two. Terms matching a pre-specified keyword list of non-central-nervous-system tissue and developmental-morphology processes, which are not interpretable in cortical tissue, were removed. *p* values were adjusted by the Benjamini-Hochberg method within each library across the retained terms, and terms at adjusted *p* < 0.05 reported. For the distance analysis, direction is expressed in proximity terms, so that a positive signed odds ratio denotes overrepresentation among genes higher near tangle-bearing neurons. For the tangle-associated trajectory analysis, a positive signed odds ratio denotes overrepresentation among genes with increasing expression towards the tangle-rich end of the trajectory.

### Reactome module selection

Reactome gene sets branching from the ‘Programmed Cell Death’, ‘Autophagy’, and ‘Cellular responses to stimuli’ root nodes were filtered by the following criteria: (1) root nodes excluded; (2) 25 ≤ number of genes in CosMx panel ≤ 80; (3) panel coverage ≥ 70% of the Reactome gene set; (4) pairwise Jaccard index ≤ 0.8; (5) one set per redundancy cluster. This left 6 programmed cell death, 4 autophagy and 10 cellular responses to stimuli modules to be scored.

### Module scoring

Modules were scored per cell with Seurat AddModuleScore on the SCT data slot, using 100 control genes per gene and 24 expression bins under a fixed seed. All modules for a given cell type were scored in a single call, and scoring was performed on the full cell type population before any tangle status or distance filter, so that the expression-matched control background is common to every analysis reported. Two models were fitted per module by linear mixed model with donor as random intercept. Proximity was tested in tangle-free cells within 1,000 μm of the nearest tangle-bearing neuron as score ∼ scaled log-distance + log_2_ total counts + negative-probe percentage + sex + scaled age + scaled *post-mortem* interval + (1|donor), with distance transformed as in other analyses. The tangle-bearing contrast was tested in all cells of the type, without a distance filter, as score ∼ tangle status + the same covariates + (1|donor). *P*values were adjusted by the Benjamini-Hochberg method across the modules of each Reactome cluster within each cell type and model, and modules at adjusted *p* < 0.05 taken as significant. Effects are reported as the change in module score per standard deviation of log-distance with its 95% confidence interval, and standardised Cohen’s *d* against the summed random-effect and residual variance. Figures show 75 μm unadjusted rolling mean with 95% confidence intervals; inference is from the mixed models, which account for the clustering of cells within donors.

Exc-IT-L2-3-*CBLN2*-*HOPX* neurons were scored for the published gene set associated with AT8-positivity,^26^ taken as genes upregulated in AT8-positive relative to AT8-negative neurons of the layer 2-3 excitatory clusters Ex1 and Ex2 at adjusted *p* < 0.05 and log_2_FC > 0.2, and intersected with the panel. Astrocytes were scored for seven published astrocyte subclusters^34^ and microglia for eight microglial states^35^.

### PHF1 intensity adjustment and sensitivity analyses

Post-run immunofluorescence provided a continuous PHF1 measurement for every cell. The 95^th^ percentile of PHF1 signal within each segmented cell mask was taken and converted to a robust *z* score within sample, centred on the median and scaled by the median absolute deviation across all cells of that sample. Cells whose mask was less than half inside the registered PHF1 image were set to missing. Because this outcome is image-derived rather than transcriptional, it was modelled with the image covariate set as intensity ∼ scaled log-distance + sex + age + *post-mortem* interval + (1|donor), in tangle-free cells within 1,000 μm.

To ask whether the transcriptional gradients were carried by this subthreshold signal, the module-score distance models were refitted with per-cell intensity added as a covariate. Collinearity was quantified by the distance-intensity correlation and by the variance inflation factor of the adjusted fit, and the distance and intensity contributions were partitioned by semi-partial R^2^, with the shared component reported as unassignable. For display, the adjusted score is the observed score minus the fitted linear contribution of intensity.

Three sensitivity analyses were performed: intensity entered instead as a within-donor percentile rank, a natural spline with three degrees of freedom, or a top-decile indicator; the distance slope refitted within each within-donor decile of intensity; and refitted after excluding all cells within 50, 100, 150 and 200 μm of a tangle-bearing neuron. Intensity is a noisy proxy for phosphorylated tau, especially after CosMx processing, and does not account for non-somatic phosphorylated tau, so a surviving slope cannot be attributed in full to a tau-independent mechanism.

### Stably expressed gene null

To test whether the reported gradients are specific to the signatures rather than a generic property of scoring a gene set on these cells, a cell-type-matched module of stably expressed genes was scored on the same axis. Stably expressed genes were identified with scMerge::scSEGIndex fitted on our own snRNA-seq reference. Reference subclusters were pooled into CosMx cell types using the mapping applied for marker-based annotation, and the reference was restricted to the CosMx panel before fitting. Size factors were computed from full-transcriptome library sizes before gene subsetting; the index was fitted within cell type, and genes above the 80^th^ percentile of the stability index were retained. Each set was scored and modelled exactly as the reported modules, with the same scoring parameters, covariates, distance transform and cap.

### CosMx nuclear morphology and rRNA stain intensity

Nucleus area was taken from the AtoMx nuclear segmentation and converted to μm^2^ using the per-cell pixel scale (1 pixel × 0.120281 = 1 μm; 1 pixel^2^ × 0.014467506 = 1 μm^2^); nuclear measures were restricted to cells with a nucleus of at least 10 μm^2^. rRNA stain intensity was the mean of the CosMx rRNA channel over each cell’s segmentation mask, log-transformed and z-scored within donor. Both were modelled at the cell level as ∼ predictor + sex + age + *post-mortem* interval + (1|donor), against distance in tangle-free neurons within 1,000 μm and against tangle status in all cells of the subtype, with BH adjustment within each measure family. Donor-paired effect sizes are reported alongside.

### IMC distance measurement analysis

For every cell the Euclidean distance to the nearest PHF1_high_ neuron was computed within the same ROI. Distances were capped at 300 μm and entered models as the plain natural logarithm divided by its s.d. and not mean-centred, with the standard deviation recomputed within each analysed cell set. Coefficients are interpretable as the change in outcome per one s.d. decrease in log-distance (“per s.d. closer”).

### IMC statistical modelling

Cell-level outcomes were modelled by linear mixed models fitted by REML (lmerTest::lmer) as outcome ∼ predictor + sex + scaled age + scaled PMI + (1 | donor).

### IMC state markers in PHF1_high_ versus PHF1_low_ neurons

Models were fitted in all excitatory neurons, with subcluster identity as an additional covariate. Ten state markers were modelled with PHF1_high_ status as the predictor, reported as log_2_FC: AT8, Aβ (4G8, MOAB-2, H31L21), BNIP3, ubiquitin FK2 and K48, LC3A/B, p62, and LAMP1.

### IMC protein gradients around tangles

The same ten markers were modelled against scaled log-distance in PHF1_low_ excitatory neurons with the same subcluster-adjusted specification and reported as log_2_FC per s.d. closer.

### IMC neuronal loss by cluster

Columnar density, the number of cells per mm of ROI width over the full grey matter depth, was averaged over each donor’s three ROIs. Loss was 100 × (mean AD / mean control density – 1). Proximity was the geometric-mean distance (uncapped) from each cluster’s PHF1_low_ cells to the nearest PHF1_high_ neuron in the same ROI. PHF1 intensity was z-scored within donor over all neuronal cells. Associations across clusters were tested by Spearman ρ, with 95% CIs from a donor bootstrap (9,999 replicates; AD and control resampled separately) and *p* values from 199,999 permutations of cluster labels.

### IMC neuronal loss by cortical depth

Each ROI (layer 1 at the top) was divided into ten equal relative-depth bands from the pia, computed over all cells. Excitatory and inhibitory columnar density were compared per band as above. PHF1 intensity was z-scored within donor over all neurons and averaged per band across AD donors. Testing was as for clusters, with permutation of band labels.

### IMC nuclear morphology

Nucleus area and eccentricity, derived from the Steinbock output which used nuclear segmentation masks, were contrasted between PHF1_high_ and PHF1_low_ neurons with a two-sided Wilcoxon signed-rank test on per-donor means, and for distance effects using the cell-level model above.

### IMC DNA quantification

DNA density is arcsinh(mean ^191^Ir ion counts within the mask) and measures DNA per unit area. Total DNA signal is the mean ^191^Ir ion count multiplied by mask area, which is the summed ^191^Ir ion count over the nucleus, on the raw multiplicative scale. This was modelled as for nucleus area.

### IMC astrocyte GFAP intensity over distance

GFAP signal intensity was z-scored within donor before fitting, so the slope is in within-donor s.d. units. Models were fitted as above.

### IMC covariate sensitivity analysis

Because ROI geometry means distance to a tangle-bearing neuron is not independent of laminar position, and because amyloid pathology is spatially structured, every IMC estimate was refitted under four nested linear mixed models (Figure S6): base; base + relative cortical depth; base + amyloid proximity; base + both. Amyloid proximity was the binary 40 μm dilated 4G8 mask term from the published object. Relative cortical depth was computed within each ROI with the following formula and z-scored:

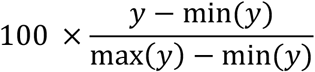

The bias in each PHF1_high_-distance estimate attributable to omitting amyloid was decomposed as γ × δ, where γ is the amyloid coefficient from the base + amyloid proximity model and δ is the coupling between amyloid and tangle proximity. δ was estimated by regressing the binary amyloid term on the outcome model’s right-hand side with the amyloid term removed. Because amyloid proximity enters the outcome models under an identity link, δ was estimated on the probability scale; a logistic fit of the same specification is reported as an odds ratio but overstates the multiplier by 1/*p*(1 – *p*). Each auxiliary fit was weighted to match its outcome model’s variance structure, so that γ × δ reproduces the observed change in the tangle coefficient. The plaque-proximal fraction within 50 μm and beyond 200 μm of a PHF1_high_ neuron was computed within donor and averaged across donors. Effect modification was tested by refitting each model with a distance × amyloid-proximity interaction and reporting the distance slope within each amyloid stratum, with BH correction applied within panel. Because the amyloid term is a 40 μm threshold on a continuous field, γ × δ is a lower bound on the bias from true amyloid proximity.

### Statistics and reproducibility

Statistical tests for individual analyses are described in the relevant methods subsection. The Benjamini-Hochberg procedure was applied to each analysis family, with significance at BH-adjusted *p* < 0.05 except for the differential gene expression analyses in which genes passed on to pathway overrepresentation analysis were filtered at adjusted *p* < 0.1. Effect sizes are reported with confidence intervals. Sample preparation, data collection, and tangle annotation were performed blinded to donor characteristics. All samples were processed and analysed identically.

## Supplemental Figures

**Figure S1.**
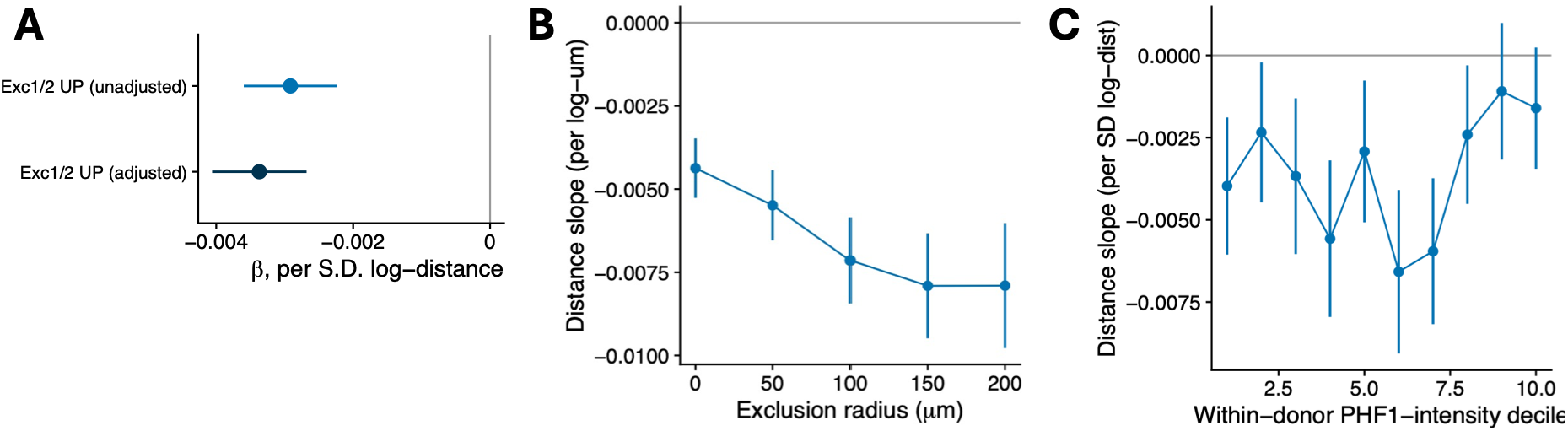
The distance gradient in tangle-free Exc-IT-L2-3-*CBLN2*-*HOPX* neurons is not explained by PHF1 intensity. Cells, signatures, covariates and mixed models as in Figure 3B (*n* = 24,657 tangle-free neurons, 9 donors); points and error bars are estimate ± 95% CI from two-sided Wald tests. **(A)** Distance slope (β per s.d. log-distance) for the reported gene set associated with AT8-positivity^26^ before and after per-cell PHF1 intensity adjustment. **(B)** Intensity-adjusted distance slope after excluding all cells within 50–200 μm of a tangle-bearing neuron. β per s.d. log-distance; *n* = 24,657 cells reduced to 16,130 after exclusion. Slopes steepen rather than attenuate: β per unit natural-log distance = -0.0044 (no exclusion; equivalent to -0.0034 per s.d. log-distance) to -0.0079; all *p* < 1.9 × 10^-16^. **(C)** Distance slope refit within each within-donor decile of per-cell PHF1 intensity (∼2,466 cells per decile; unadjusted), for the reported signature. All ten deciles are negative, including the dimmest decile (β = -0.0040 [-0.0061, -0.0019], *p* = 1.9 × 10^-4^).

**Figure S2.**
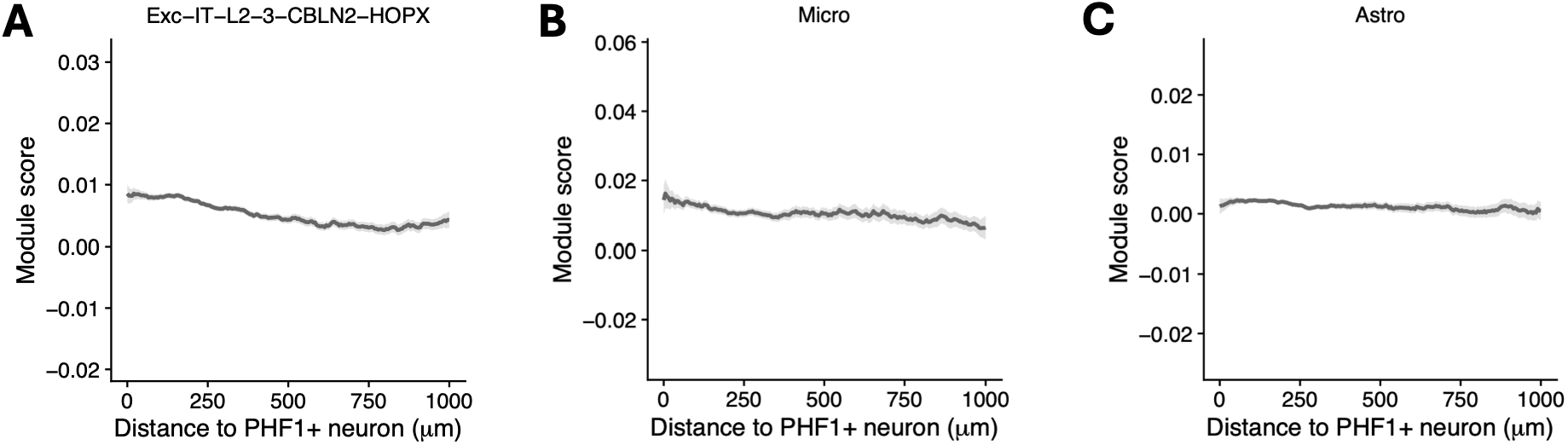
Stably expressed genes show no comparable spatial gradient. Rolling mean (75 μm window) of a cell-type-specific stably expressed gene module (scSEGIndex, derived from our snRNA- seq reference) over distance to the nearest tangle-bearing neuron, in tangle-free cells. Shading, 95% CI; 9 donors. Each *y*-axis spans the same module-score range as the corresponding panel; absolute offsets differ because each module is centred on its own expression-matched controls. **(A)** Exc-IT-L2-3-*CBLN2*-*HOPX* neurons (305 genes; 24,672 tangle-free neurons; *y*-axis span matches Figure 3B). β = -3.6 × 10^-4^ per s.d. log-distance (95% CI -5.3 × 10^-4^ to -1.9 × 10^-4^; adjusted *p* = 2.1 × 10^-4^). The slope is only 10.7% of the reported effect in Figure 3B. **(B)** Microglia (237 genes; 6,524 cells; Figure 6B). β = -1.7 × 10^-4^ per s.d. log-distance (95% CI -6.0 × 10^-4^ to +2.6 × 10^-4^; adjusted *p* = 0.52). **(C)** Astrocytes (284 genes, 16,552 cells; Figure 6C). β = -5.9 × 10^-5^ per s.d. log-distance (95% CI -2.4 × 10^-4^ to +1.2 × 10^-4^; adjusted *p* = 0.52).

**Figure S3.**
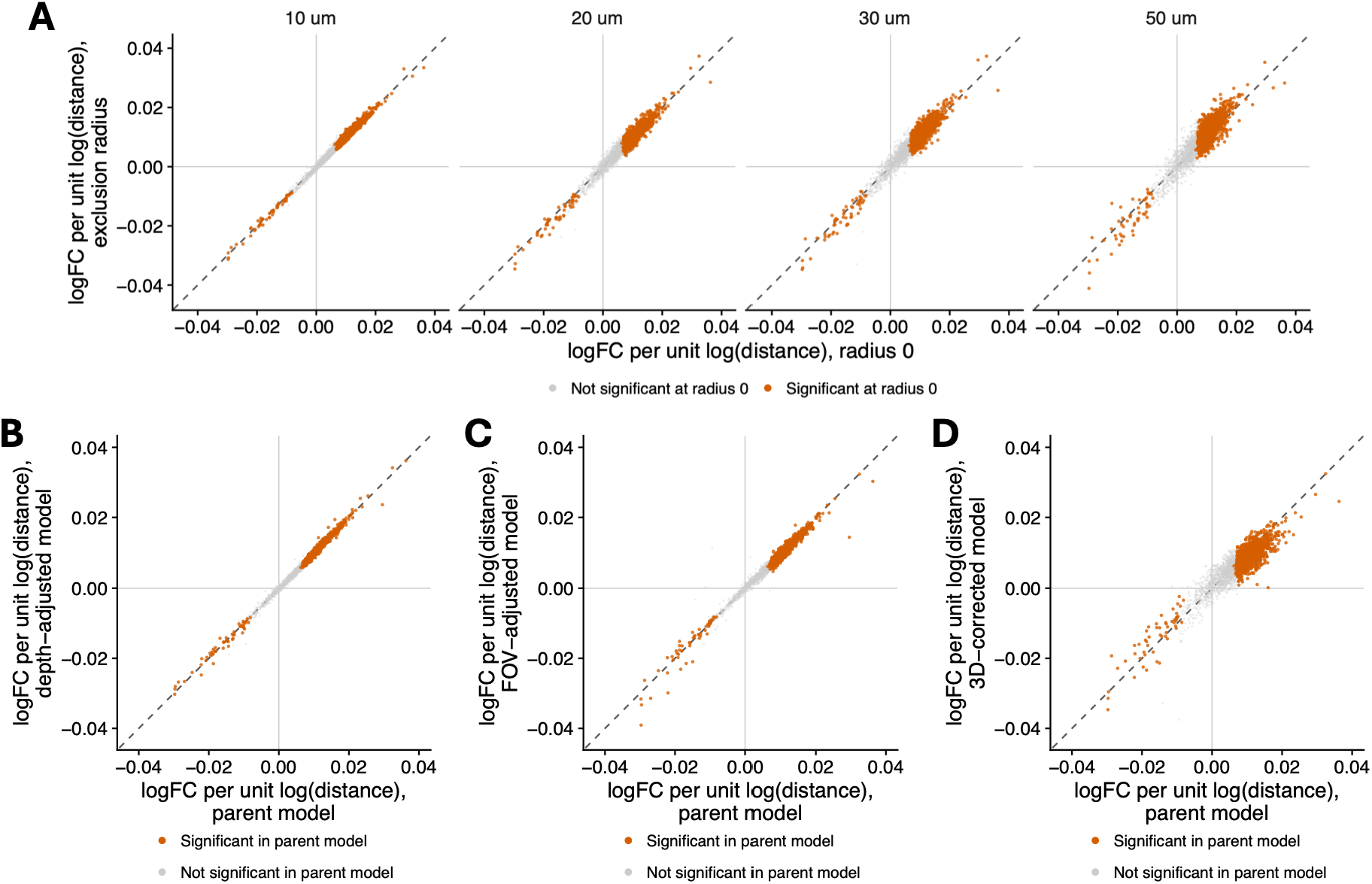
Robustness of the tangle-proximity transcriptional gradients. **(A–D)** Per-gene distance coefficients (log_2_FC per s.d. log μm; cell-level linear mixed model in tangle-free Exc-IT-L2-3- *CBLN2*-*HOPX* neurons) from the primary model (*x* axis) versus: models refit after excluding cells within 10, 20, 30 or 50 μm of the nearest tangle-bearing neuron (**A**; *y* axis; 0.1–3.1% of cells excluded); model refit after the addition of scaled cortical depth as a covariate (**B**); after the addition of a random intercept for imaging field of view (517 FOVs nested within donor), so that the distance coefficient is estimated from within-FOV variation only (**C**); and after the simulation of tangle location in the third dimension, outside of the observable field of view, with recalculation of distance to nearest tangle and refitting of the model (**D**). All 2,468 tested genes are shown; orange, the 1,110 genes significant in the primary model (BH-adjusted *p* < 0.1); grey, all others. Dashed line, identity. Gene set and axis limits are identical across panels. No significant gene changes sign in any sensitivity contrast; **A**, Spearman ρ = 0.89–0.99 (Table S8); **B**, ρ = 0.995; **C**, ρ = 0.99 ; **D**, ρ = 0.86. All *p* < 10^-300^.

**Figure S4.**
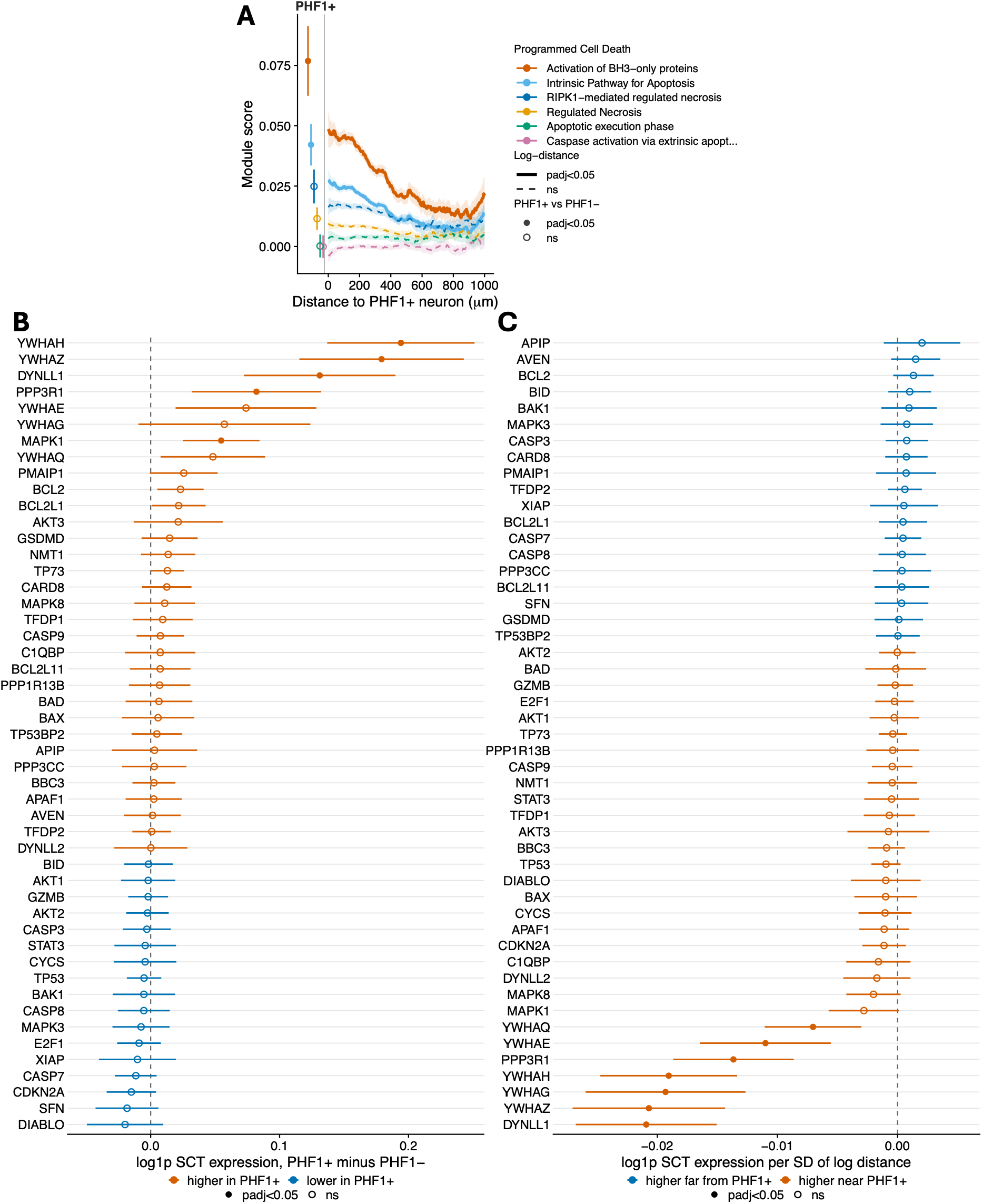
Apparent cell death module enrichment in and around tangle-bearing neurons is driven by genes encoding the anti-apoptotic 14-3-3 scaffold, not by apoptotic effectors. **(A)** Gene sets from the ‘Programmed cell death’ Reactome cluster were filtered (see STAR Methods) and scored (AddModuleScore) in Exc-IT-L2-3-*CBLN2*-*HOPX* neurons as in Figures 3F **and 3G**. Scores were elevated in tangle-free neurons with increasing proximity to tangles and in tangle-bearing neurons for activation of BH3-only proteins (tangle-bearing, adjusted *p* = 1.2 × 10^-7^; proximity, adjusted *p* = 1.0 × 10^-14^) and intrinsic pathway for apoptosis (tangle-bearing, adjusted *p* = 1.1 × 10^-6^; proximity, adjusted *p* = 1.7 × 10^-11^) gene sets. **(B and C)** Per-gene decomposition of the intrinsic pathway for apoptosis module score in Exc-IT-L2-3-*CBLN2*-*HOPX* neurons, for the tangle-status (**B**) and proximity (**C**) contrasts. Points are linear mixed-model coefficients with 95% CI from ∼ predictor + nUMI + negative probe fraction + sex + age + PMI + (1|donor); genes ordered by effect size; filled points, BH- adjusted *p* < 0.05 within the module. **B**, Tangle-bearing minus tangle-free neurons; *n* = 26,585 neurons (188 tangle-bearing), 9 donors; 5 of 49 genes significant, 4 of which (*YWHAH*, *YWHAZ*, *DYNLL1*, *MAPK1*) encode proteins that restrain rather than execute apoptosis. **C**, Per s.d. of log-distance to the nearest tangle-bearing neuron; tangle-free neurons within 1,000 μm; *n* = 24,672 neurons, 9 donors; 7 of 49 genes significant, 6 of them 14-3-3 paralogues or *DYNLL1*. In neither contrast is an executioner caspase, *BAX* or *BAK1*, or an apoptosome component (*APAF1*, *CYCS*) significant.

**Figure S5.**
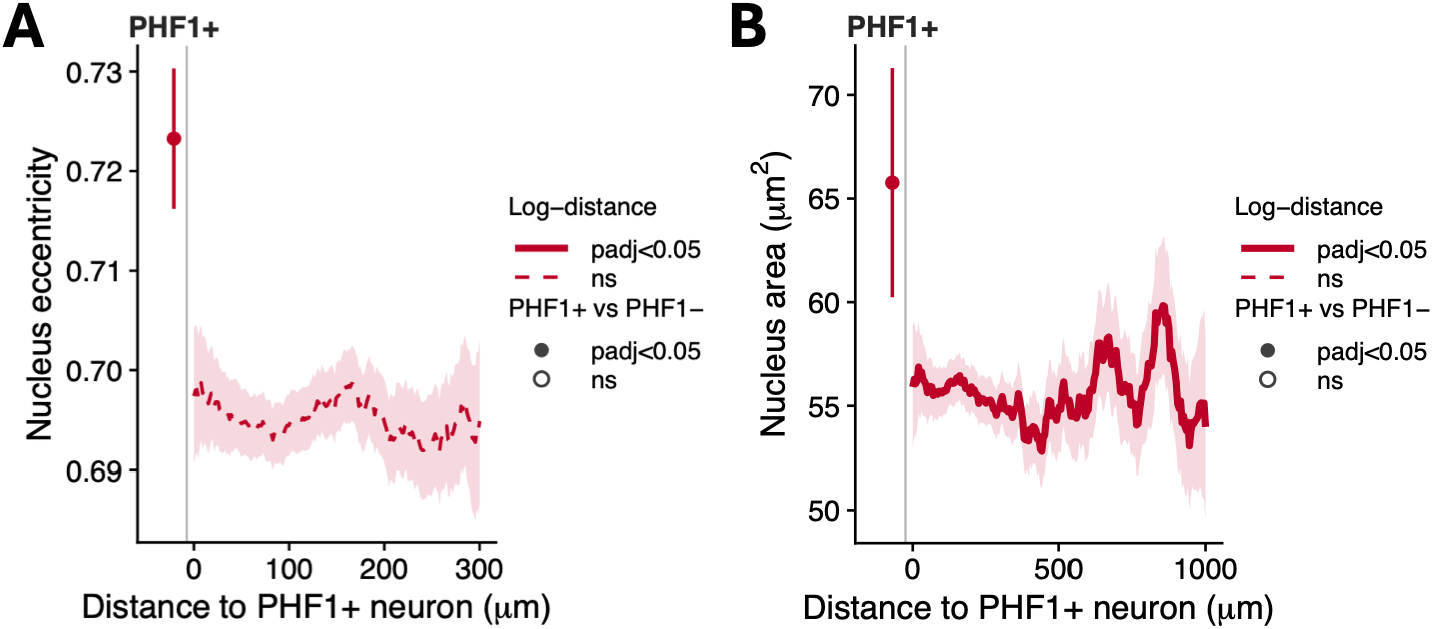
Nucleus eccentricity and CosMx replication of nuclear enlargement. Cells, models, curves and encodings as in Figures 5A–C. **(A)** Nucleus eccentricity in excitatory neurons (IMC). PHF1_high_ nuclei are more eccentric (+0.023, 95% CI 0.012–0.035, +3.4%, *d_z_* = 0.64, 34/44 donors concordant, adjusted *p* = 1.8 × 10^-4^; filled point). **(B)** Nucleus area in Exc-IT-L2-3-CBLN2-HOPX neurons (CosMx). Tangle-bearing nuclei are 20.0% larger (+11.1 μm^2^, 95% CI 6.0–16.2, *d_z_* = 0.86, 7/9 donors concordant, adjusted *p* = 4.9 × 10^-5^, *n* = 18,627 cells, *n* = 9 donors; cell-level mixed model), reproducing Figure 5A in an independent modality (1 sample was in both the CosMx and AD IMC cohorts). Distance gradient: β = +1.11 μm^2^ per s.d. closer (adjusted *p* = 1.3 × 10^-4^). Nuclei ≥ 10 μm^2^ only.

**Figure S6.**
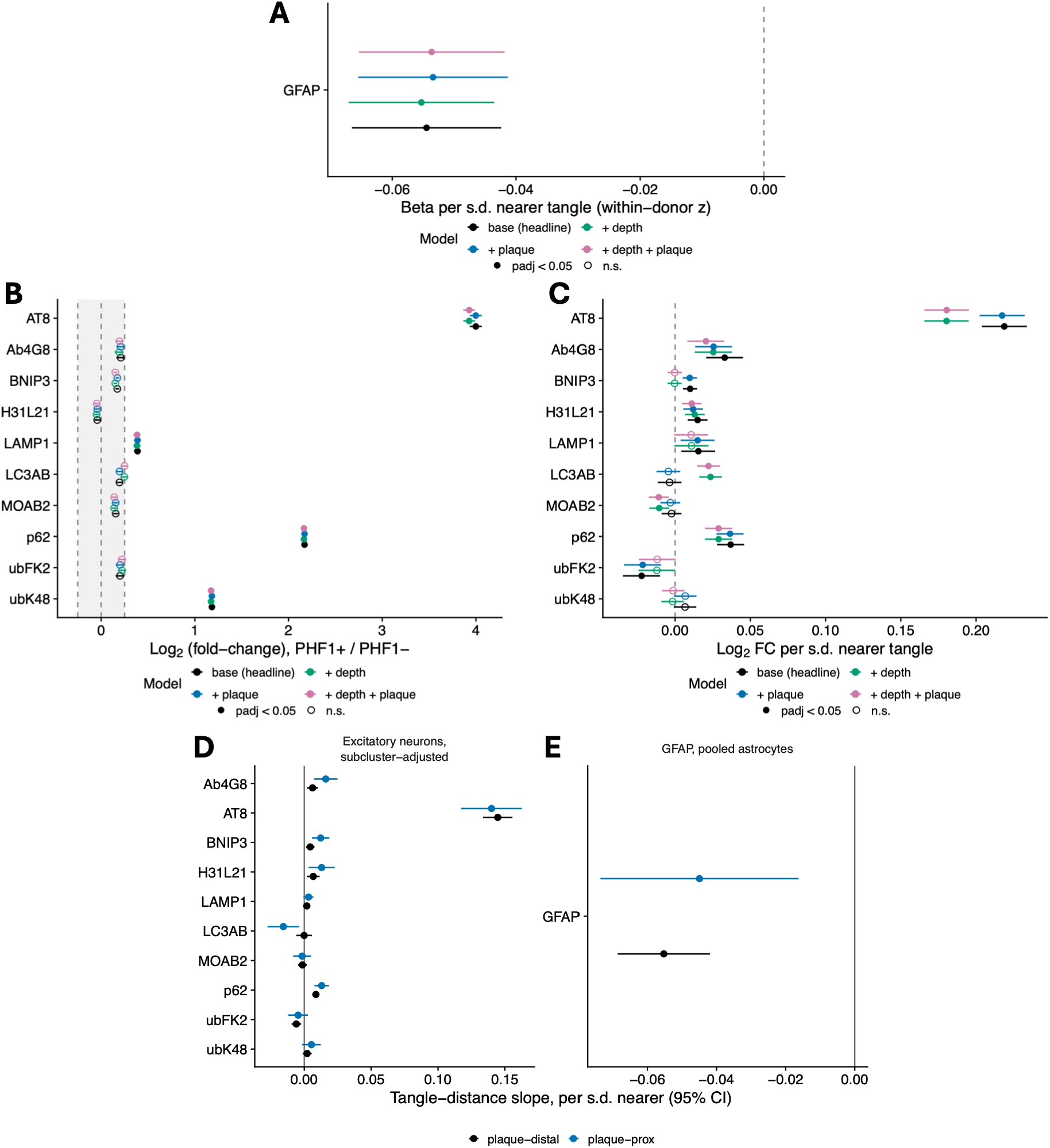
Adjustment for Aβ plaque proximity and cortical depth. Predictor estimates (dots) with 95% CI (bars) in IMC data under four nested linear mixed models: base (as used in the main figures), + depth, + amyloid, and + both. **(A)** GFAP in pooled astrocytes (within-donor *z*) versus distance to the nearest PHF1_high_ neuron (Figure 6D; 26,832 cells). **(B)** IMC state markers in PHF1_high_ versus PHF1_low_ excitatory neurons (Figure 2H; 36,133 cells); shading, |log_2_FC| < 0.25. **(C)** As in **B** versus proximity to PHF1_high_ neuron (Figure 3H; 26,653 cells). *n* = 44 AD donors. Models: outcome ∼ predictor + subcluster (**B** and **C**) + sex + age + PMI + (1|donor), REML, two-sided. Amyloid, nucleus within a 40 μm dilated 4G8 mask; depth, relative cortical depth per s.d. (0 = pia). Filled dots, adjusted *p* < 0.05 (**B**, additionally |log_2_FC| ≥ 0.25). Adjustment leaves **A** and **B** essentially unchanged (median |Δβ| 1.5% and 8.5%; no sign changes). Five markers in **C** cross the significance threshold (BNIP3, LAMP1 and ubFK2 lose significance; LC3A/B and MOAB-2 gain significance), consistent with their proximity to the threshold in the base model. **(D)** As in **C**, stratified by plaque proximity: PHF1_high_-distance slopes (per s.d. nearer; 95% CI) in plaque-distal (black) and plaque-proximal (blue) excitatory neurons (26,653 cells). **(E)** As in **A**, stratified likewise (26,832 cells). Slopes are from one model per marker with an added distance × plaque-proximity interaction, otherwise as above. No interaction survived FDR correction (0/11), and plaque-distal slopes matched the base estimates (AT8, 0.145 vs 0.145; GFAP, -0.055 vs -0.054).

**Figure S7.**
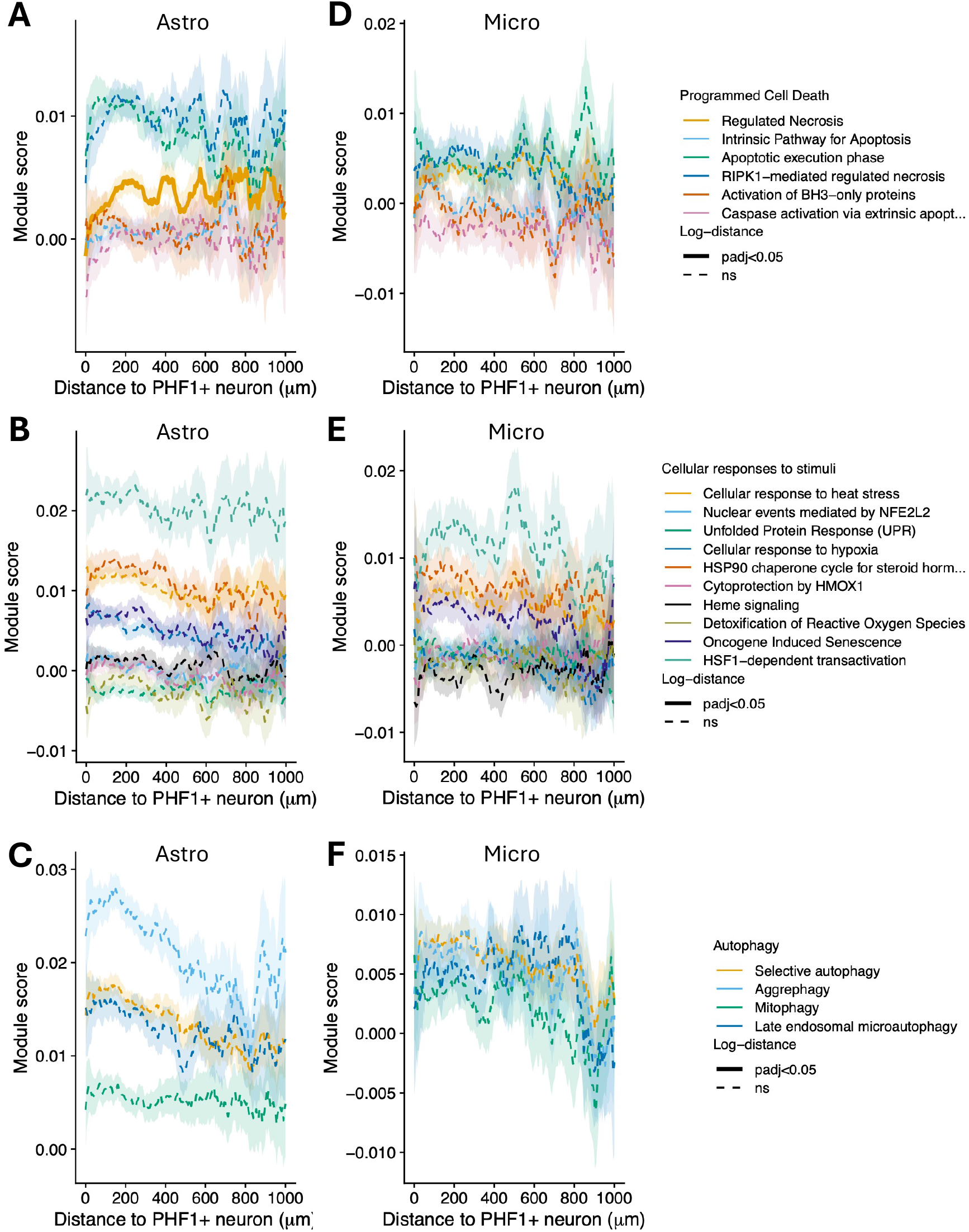
Cell death, stress and autophagy modules in glia did not vary detectably with tangle proximity. Reactome modules of the programmed cell death (**A** and **D**), cellular responses to stimuli (**B** and **E**) and autophagy (**C** and **F**) clusters scored in astrocytes (**A–C**; *n* = 16,552 cells, 9 donors) and microglia (**D–F**; *n* = 6,524 cells, 9 donors) against distance to the nearest tangle-bearing neuron. Module filtering, scoring, models and line encodings as in Figures 3F**, 3G** and **S4A**. Only regulated necrosis in astrocytes had a significant log-distance term (β = 7.7 × 10^-4^ per s.d. log-distance, 95% CI 3.4 × 10^-4^ to 1.2 × 10^-3^, adjusted *p* = 0.0026; lower near tangles); all other modules, adjusted *p* > 0.05.

## References

1. Kidd, M. (1963). Paired helical filaments in electron microscopy of Alzheimer’s disease. Nature 197, 192–193. 10.1038/197192b0.

2. Bramblett, G.T., Goedert, M., Jakes, R., Merrick, S.E., Trojanowski, J.Q., and Lee, V.M. (1993). Abnormal tau phosphorylation at Ser396 in Alzheimer’s disease recapitulates development and contributes to reduced microtubule binding. Neuron 10, 1089–1099. 10.1016/0896-6273(93)90057-x.

3. Li, X., Kumar, Y., Zempel, H., Mandelkow, E.-M., Biernat, J., and Mandelkow, E. (2011). Novel diffusion barrier for axonal retention of Tau in neurons and its failure in neurodegeneration. EMBO J. 30, 4825–4837. 10.1038/emboj.2011.376.

4. Shi, Y., Zhang, W., Yang, Y., Murzin, A.G., Falcon, B., Kotecha, A., van Beers, M., Tarutani, A., Kametani, F., Garringer, H.J., et al. (2021). Structure-based classification of tauopathies. Nature 598, 359–363. 10.1038/s41586-021-03911-7.

5. Fitzpatrick, A.W.P., Falcon, B., He, S., Murzin, A.G., Murshudov, G., Garringer, H.J., Crowther, R.A., Ghetti, B., Goedert, M., and Scheres, S.H.W. (2017). Cryo-EM structures of tau filaments from Alzheimer’s disease. Nature 547, 185–190. 10.1038/nature23002.

6. Arendt, T., Stieler, J., Strijkstra, A.M., Hut, R.A., Rüdiger, J., Van Der Zee, E.A., Harkany, T., Holzer, M., and Härtig, W. (2003). Reversible Paired Helical Filament-Like Phosphorylation of Tau Is an Adaptive Process Associated with Neuronal Plasticity in Hibernating Animals. J. Neurosci. 23, 6972–6981. 10.1523/JNEUROSCI.23-18-06972.2003.

7. Stieler, J.T., Bullmann, T., Kohl, F., Tøien, Ø., Brückner, M.K., Härtig, W., Barnes, B.M., and Arendt, T. (2011). The physiological link between metabolic rate depression and tau phosphorylation in mammalian hibernation. PloS One 6, e14530. 10.1371/journal.pone.0014530.

8. Planel, E., Richter, K.E.G., Nolan, C.E., Finley, J.E., Liu, L., Wen, Y., Krishnamurthy, P., Herman, M., Wang, L., Schachter, J.B., et al. (2007). Anesthesia leads to tau hyperphosphorylation through inhibition of phosphatase activity by hypothermia. J. Neurosci. Off. J. Soc. Neurosci. 27, 3090–3097. 10.1523/JNEUROSCI.4854-06.2007.

9. Kanemaru, K., Takio, K., Miura, R., Titani, K., and Ihara, Y. (1992). Fetal-type phosphorylation of the tau in paired helical filaments. J. Neurochem. 58, 1667–1675. 10.1111/j.1471-4159.1992.tb10039.x.

10. Kenessey, A., and Yen, S.H. (1993). The extent of phosphorylation of fetal tau is comparable to that of PHF-tau from Alzheimer paired helical filaments. Brain Res. 629, 40–46. 10.1016/0006-8993(93)90478-6.

11. Kimura, T., Whitcomb, D.J., Jo, J., Regan, P., Piers, T., Heo, S., Brown, C., Hashikawa, T., Murayama, M., Seok, H., et al. (2014). Microtubule-associated protein tau is essential for long-term depression in the hippocampus. Philos. Trans. R. Soc. Lond. B. Biol. Sci. *369*, 20130144. 10.1098/rstb.2013.0144.

12. Ittner, A., Chua, S.W., Bertz, J., Volkerling, A., van der Hoven, J., Gladbach, A., Przybyla, M., Bi, M., van Hummel, A., Stevens, C.H., et al. (2016). Site-specific phosphorylation of tau inhibits amyloid-β toxicity in Alzheimer’s mice. Science 354, 904–908. 10.1126/science.aah6205.

13. Gong, C.X., Singh, T.J., Grundke-Iqbal, I., and Iqbal, K. (1993). Phosphoprotein phosphatase activities in Alzheimer disease brain. J. Neurochem. 61, 921–927. 10.1111/j.1471-4159.1993.tb03603.x.

14. Gong, C.X., Shaikh, S., Wang, J.Z., Zaidi, T., Grundke-Iqbal, I., and Iqbal, K. (1995). Phosphatase activity toward abnormally phosphorylated tau: decrease in Alzheimer disease brain. J. Neurochem. 65, 732–738. 10.1046/j.1471-4159.1995.65020732.x.

15. Sontag, E., Hladik, C., Montgomery, L., Luangpirom, A., Mudrak, I., Ogris, E., and White, C.L. (2004). Downregulation of protein phosphatase 2A carboxyl methylation and methyltransferase may contribute to Alzheimer disease pathogenesis. J. Neuropathol. Exp. Neurol. 63, 1080–1091. 10.1093/jnen/63.10.1080.

16. Patrick, G.N., Zukerberg, L., Nikolic, M., de la Monte, S., Dikkes, P., and Tsai, L.H. (1999). Conversion of p35 to p25 deregulates Cdk5 activity and promotes neurodegeneration. Nature 402, 615–622. 10.1038/45159.

17. Pei, J.J., Tanaka, T., Tung, Y.C., Braak, E., Iqbal, K., and Grundke-Iqbal, I. (1997). Distribution, levels, and activity of glycogen synthase kinase-3 in the Alzheimer disease brain. J. Neuropathol. Exp. Neurol. 56, 70–78. 10.1097/00005072-199701000-00007.

18. Arriagada, P.V., Growdon, J.H., Hedley-Whyte, E.T., and Hyman, B.T. (1992). Neurofibrillary tangles but not senile plaques parallel duration and severity of Alzheimer’s disease. Neurology 42, 631–639. 10.1212/wnl.42.3.631.

19. Gómez-Isla, T., Hollister, R., West, H., Mui, S., Growdon, J.H., Petersen, R.C., Parisi, J.E., and Hyman, B.T. (1997). Neuronal loss correlates with but exceeds neurofibrillary tangles in Alzheimer’s disease. Ann. Neurol. 41, 17–24. 10.1002/ana.410410106.

20. Kril, J.J., Patel, S., Harding, A.J., and Halliday, G.M. (2002). Neuron loss from the hippocampus of Alzheimer’s disease exceeds extracellular neurofibrillary tangle formation. Acta Neuropathol. (Berl.) 103, 370–376. 10.1007/s00401-001-0477-5.

21. de Calignon, A., Fox, L.M., Pitstick, R., Carlson, G.A., Bacskai, B.J., Spires-Jones, T.L., and Hyman, B.T. (2010). Caspase activation precedes and leads to tangles. Nature 464, 1201–1204. 10.1038/nature08890.

22. Kuchibhotla, K.V., Wegmann, S., Kopeikina, K.J., Hawkes, J., Rudinskiy, N., Andermann, M.L., Spires-Jones, T.L., Bacskai, B.J., and Hyman, B.T. (2014). Neurofibrillary tangle-bearing neurons are functionally integrated in cortical circuits in vivo. Proc. Natl. Acad. Sci. U. S. A. 111, 510–514. 10.1073/pnas.1318807111.

23. Zwang, T.J., Sastre, E.D., Wolf, N., Ruiz-Uribe, N., Woost, B., Hoglund, Z., Fan, Z., Bailey, J., Nfor, L., Buée, L., et al. (2024). Neurofibrillary tangle-bearing neurons have reduced risk of cell death in mice with Alzheimer’s pathology. Cell Rep. 43, 114574. 10.1016/j.celrep.2024.114574.

24. Morsch, R., Simon, W., and Coleman, P.D. (1999). Neurons may live for decades with neurofibrillary tangles. J. Neuropathol. Exp. Neurol. 58, 188–197. 10.1097/00005072-199902000-00008.

25. Spires-Jones, T.L., Calignon, A. de, Matsui, T., Zehr, C., Pitstick, R., Wu, H.-Y., Osetek, J.D., Jones, P.B., Bacskai, B.J., Feany, M.B., et al. (2008). In Vivo Imaging Reveals Dissociation between Caspase Activation and Acute Neuronal Death in Tangle-Bearing Neurons. J. Neurosci. 28, 862–867. 10.1523/JNEUROSCI.3072-08.2008.

26. Otero-Garcia, M., Mahajani, S.U., Wakhloo, D., Tang, W., Xue, Y.-Q., Morabito, S., Pan, J., Oberhauser, J., Madira, A.E., Shakouri, T., et al. (2022). Molecular signatures underlying neurofibrillary tangle susceptibility in Alzheimer’s disease. Neuron 110, 2929–2948.e8. 10.1016/j.neuron.2022.06.021.

27. Foiani, M.S., Bourdenx, M., Kraller, L., Nirujogi, R.S., Yiu, A., Davies, H., Patel, S., Damoc, L.S., Mitchener, L., Jaunmuktane, Z., et al. (2026). Spatial Single-Cell Proteomics Reveals Molecular Trajectories Of Tangle-Bearing Neurons In Alzheimer’s Disease. Preprint at bioRxiv, 10.64898/2026.04.26.720932 10.64898/2026.04.26.720932.

28. Boulger, S.L., Lerose, D., Adair, E., Papageorgopoulou, M., Zhao, Y., Avot, B., Thomas, M., Nibber, P., Klug, H., Fancy, N.N., et al. (2026). Intrinsic molecular susceptibility underlies selective neuronal vulnerability in the Alzheimer’s disease entorhinal cortex. Preprint at bioRxiv, 10.64898/2026.06.22.733753 10.64898/2026.06.22.733753.

29. Mathys, H., Boix, C.A., Akay, L.A., Xia, Z., Davila-Velderrain, J., Ng, A.P., Jiang, X., Abdelhady, G., Galani, K., Mantero, J., et al. (2024). Single-cell multiregion dissection of Alzheimer’s disease. Nature 632, 858–868. 10.1038/s41586-024-07606-7.

30. McHugh, M.L. (2012). Interrater reliability: the kappa statistic. Biochem. Medica 22, 276–282.

31. Zha, J., Harada, H., Yang, E., Jockel, J., and Korsmeyer, S.J. (1996). Serine phosphorylation of death agonist BAD in response to survival factor results in binding to 14-3-3 not BCL-X(L). Cell 87, 619–628. 10.1016/s0092-8674(00)81382-3.

32. Paonessa, F., Evans, L.D., Solanki, R., Larrieu, D., Wray, S., Hardy, J., Jackson, S.P., and Livesey, F.J. (2019). Microtubules Deform the Nuclear Membrane and Disrupt Nucleocytoplasmic Transport in Tau-Mediated Frontotemporal Dementia. Cell Rep. 26, 582–593.e5. 10.1016/j.celrep.2018.12.085.

33. Frost, B., Hemberg, M., Lewis, J., and Feany, M.B. (2014). Tau promotes neurodegeneration through global chromatin relaxation. Nat. Neurosci. 17, 357–366. 10.1038/nn.3639.

34. Serrano-Pozo, A., Li, H., Li, Z., Muñoz-Castro, C., Jaisa-aad, M., Healey, M.A., Welikovitch, L.A., Jayakumar, R., Bryant, A.G., Noori, A., et al. (2024). Astrocyte transcriptomic changes along the spatiotemporal progression of Alzheimer’s disease. Nat. Neurosci. 27, 2384–2400. 10.1038/s41593-024-01791-4.

35. Mancuso, R., Fattorelli, N., Martinez-Muriana, A., Davis, E., Wolfs, L., Van Den Daele, J., Geric, I., Premereur, J., Polanco, P., Bijnens, B., et al. (2024). Xenografted human microglia display diverse transcriptomic states in response to Alzheimer’s disease-related amyloid-β pathology. Nat. Neurosci. 27, 886–900. 10.1038/s41593-024-01600-y.

36. Nixon, R.A., Wegiel, J., Kumar, A., Yu, W.H., Peterhoff, C., Cataldo, A., and Cuervo, A.M. (2005). Extensive involvement of autophagy in Alzheimer disease: an immuno-electron microscopy study. J. Neuropathol. Exp. Neurol. 64, 113–122. 10.1093/jnen/64.2.113.

37. Boland, B., Kumar, A., Lee, S., Platt, F.M., Wegiel, J., Yu, W.H., and Nixon, R.A. (2008). Autophagy Induction and Autophagosome Clearance in Neurons: Relationship to Autophagic Pathology in Alzheimer’s Disease. J. Neurosci. 28, 6926–6937. 10.1523/JNEUROSCI.0800-08.2008.

38. Dickey, C.A., Kamal, A., Lundgren, K., Klosak, N., Bailey, R.M., Dunmore, J., Ash, P., Shoraka, S., Zlatkovic, J., Eckman, C.B., et al. (2007). The high-affinity HSP90-CHIP complex recognizes and selectively degrades phosphorylated tau client proteins. J. Clin. Invest. 117, 648–658. 10.1172/JCI29715.

39. Escartin, C., Galea, E., Lakatos, A., O’Callaghan, J.P., Petzold, G.C., Serrano-Pozo, A., Steinhäuser, C., Volterra, A., Carmignoto, G., Agarwal, A., et al. (2021). Reactive astrocyte nomenclature, definitions, and future directions. Nat. Neurosci. 24, 312–325. 10.1038/s41593-020-00783-4.

40. Hao, Y., Stuart, T., Kowalski, M.H., Choudhary, S., Hoffman, P., Hartman, A., Srivastava, A., Molla, G., Madad, S., Fernandez-Granda, C., et al. (2024). Dictionary learning for integrative, multimodal and scalable single-cell analysis. Nat. Biotechnol. 42, 293–304. 10.1038/s41587-023-01767-y.

41. Amezquita, R.A., Lun, A.T.L., Becht, E., Carey, V.J., Carpp, L.N., Geistlinger, L., Marini, F., Rue-Albrecht, K., Risso, D., Soneson, C., et al. (2020). Orchestrating single-cell analysis with Bioconductor. Nat. Methods 17, 137–145. 10.1038/s41592-019-0654-x.

42. Ritchie, M.E., Phipson, B., Wu, D., Hu, Y., Law, C.W., Shi, W., and Smyth, G.K. (2015). limma powers differential expression analyses for RNA-sequencing and microarray studies. Nucleic Acids Res. 43, e47. 10.1093/nar/gkv007.

43. Chen, Y., Chen, L., Lun, A.T.L., Baldoni, P.L., and Smyth, G.K. (2025). edgeR v4: powerful differential analysis of sequencing data with expanded functionality and improved support for small counts and larger datasets. Nucleic Acids Res. 53, gkaf018. 10.1093/nar/gkaf018.

44. Hoffman, G.E., and Roussos, P. (2021). Dream: powerful differential expression analysis for repeated measures designs. Bioinformatics 37, 192–201. 10.1093/bioinformatics/btaa687.

45. Bates, D., Mächler, M., Bolker, B., and Walker, S. (2015). Fitting Linear Mixed-Effects Models Using lme4. J. Stat. Softw. 67, 1–48. 10.18637/jss.v067.i01.

46. Kuznetsova, A., Brockhoff, P.B., and Christensen, R.H.B. (2017). lmerTest Package: Tests in Linear Mixed Effects Models. J. Stat. Softw. 82, 1–26. 10.18637/jss.v082.i13.

47. Kuleshov, M.V., Jones, M.R., Rouillard, A.D., Fernandez, N.F., Duan, Q., Wang, Z., Koplev, S., Jenkins, S.L., Jagodnik, K.M., Lachmann, A., et al. (2016). Enrichr: a comprehensive gene set enrichment analysis web server 2016 update. Nucleic Acids Res. 44, W90–97. 10.1093/nar/gkw377.

48. Aibar, S., González-Blas, C.B., Moerman, T., Huynh-Thu, V.A., Imrichova, H., Hulselmans, G., Rambow, F., Marine, J.-C., Geurts, P., Aerts, J., et al. (2017). SCENIC: single-cell regulatory network inference and clustering. Nat. Methods 14, 1083–1086. 10.1038/nmeth.4463.

49. Gu, Z., Eils, R., and Schlesner, M. (2016). Complex heatmaps reveal patterns and correlations in multidimensional genomic data. Bioinformatics 32, 2847–2849. 10.1093/bioinformatics/btw313.

50. Zeileis, A., and Grothendieck, G. (2005). zoo: S3 Infrastructure for Regular and Irregular Time Series. J. Stat. Softw. 14, 1–27. 10.18637/jss.v014.i06.

51. Ahlmann-Eltze, C., and Huber, W. (2021). glmGamPoi: fitting Gamma-Poisson generalized linear models on single cell count data. Bioinformatics 36, 5701–5702. 10.1093/bioinformatics/btaa1009.

52. Germain, P.-L., Lun, A., Garcia Meixide, C., Macnair, W., and Robinson, M.D. (2021). Doublet identification in single-cell sequencing data using scDblFinder. F1000Research *10*, 979. 10.12688/f1000research.73600.2.

53. Lin, Y., Ghazanfar, S., Wang, K.Y.X., Gagnon-Bartsch, J.A., Lo, K.K., Su, X., Han, Z.-G., Ormerod, J.T., Speed, T.P., Yang, P., et al. (2019). scMerge leverages factor analysis, stable expression, and pseudoreplication to merge multiple single-cell RNA-seq datasets. Proc. Natl. Acad. Sci. U. S. A. 116, 9775–9784. 10.1073/pnas.1820006116.

54. McInnes, L., Healy, J., Saul, N., and Großberger, L. (2018). UMAP: Uniform Manifold Approximation and Projection. J. Open Source Softw. 3, 861. 10.21105/joss.00861.

55. Korsunsky, I., Millard, N., Fan, J., Slowikowski, K., Zhang, F., Wei, K., Baglaenko, Y., Brenner, M., Loh, P., and Raychaudhuri, S. (2019). Fast, sensitive and accurate integration of single-cell data with Harmony. Nat. Methods 16, 1289–1296. 10.1038/s41592-019-0619-0.

56. Street, K., Risso, D., Fletcher, R.B., Das, D., Ngai, J., Yosef, N., Purdom, E., and Dudoit, S. (2018). Slingshot: cell lineage and pseudotime inference for single-cell transcriptomics. BMC Genomics 19, 477. 10.1186/s12864-018-4772-0.

57. Windhager, J., Zanotelli, V.R.T., Schulz, D., Meyer, L., Daniel, M., Bodenmiller, B., and Eling, N. (2023). An end-to-end workflow for multiplexed image processing and analysis. Nat. Protoc. 18, 3565–3613. 10.1038/s41596-023-00881-0.

58. Harris, C.R., Millman, K.J., van der Walt, S.J., Gommers, R., Virtanen, P., Cournapeau, D., Wieser, E., Taylor, J., Berg, S., Smith, N.J., et al. (2020). Array programming with NumPy. Nature 585, 357–362. 10.1038/s41586-020-2649-2.

59. Walt, S. van der, Schönberger, J.L., Nunez-Iglesias, J., Boulogne, F., Warner, J.D., Yager, N., Gouillart, E., and Yu, T. (2014). scikit-image: image processing in Python. PeerJ 2, e453. 10.7717/peerj.453.

60. Milacic, M., Beavers, D., Conley, P., Gong, C., Gillespie, M., Griss, J., Haw, R., Jassal, B., Matthews, L., May, B., et al. (2024). The Reactome Pathway Knowledgebase 2024. Nucleic Acids Res. 52, D672–D678. 10.1093/nar/gkad1025.

61. He, S., Bhatt, R., Brown, C., Brown, E.A., Buhr, D.L., Chantranuvatana, K., Danaher, P., Dunaway, D., Garrison, R.G., Geiss, G., et al. (2022). High-plex imaging of RNA and proteins at subcellular resolution in fixed tissue by spatial molecular imaging. Nat. Biotechnol. 40, 1794–1806. 10.1038/s41587-022-01483-z.

62. Landis, J.R., and Koch, G.G. (1977). The measurement of observer agreement for categorical data. Biometrics 33, 159–174.

63. Gwet, K.L. (2008). Computing inter-rater reliability and its variance in the presence of high agreement. Br. J. Math. Stat. Psychol. 61, 29–48. 10.1348/000711006X126600.

