## Supplementary material for "Neuronal and glial transcriptional states vary with proximity to tangle-bearing neurons in the human entorhinal cortex": Figure S1

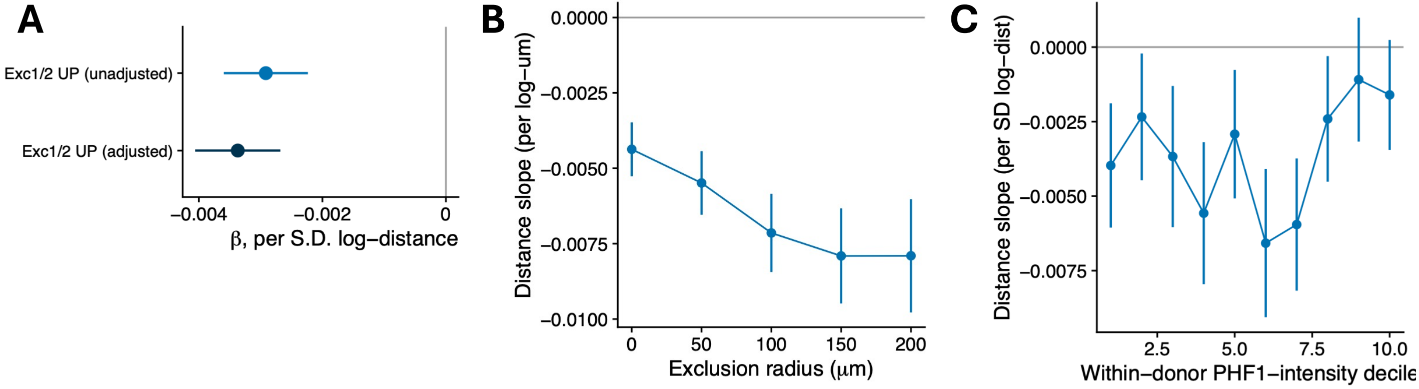

A
Exc1/2 UP (unadjusted)
Exc1/2 UP (adjusted)
−0.004
−0.002
0
β, per S.D. log−distance
B
0.0000
−0.0025
−0.0050
−0.0075
−0.0100
Distance slope (per log−um)
0
50
100
150
200
Exclusion radius (μm)
C
0.0000
−0.0025
−0.0050
−0.0075
Distance slope (per SD log−dist)
2.5
5.0
7.5
10.0
Within−donor PHF1−intensity decile
