## Supplementary material for "Neuronal and glial transcriptional states vary with proximity to tangle-bearing neurons in the human entorhinal cortex": Figure S2

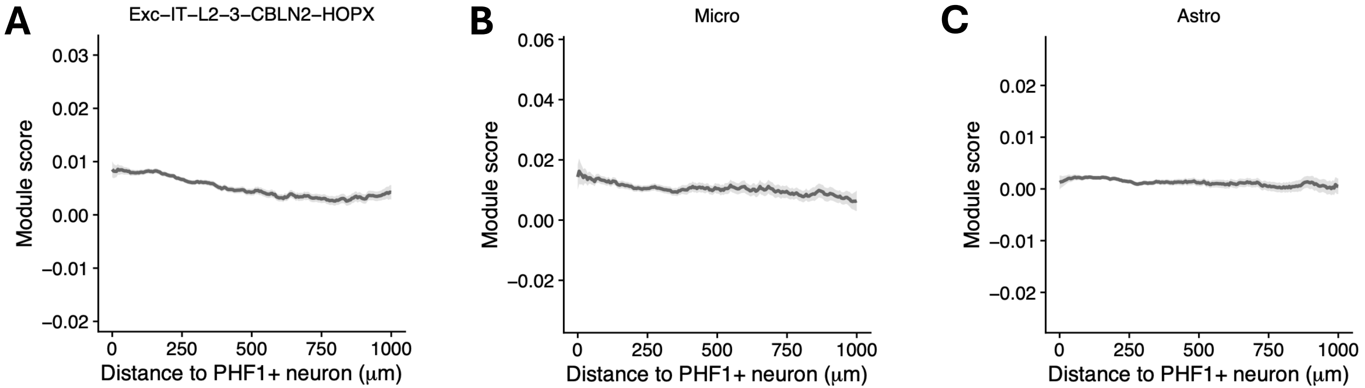
A
Exc–IT–L2–3–CBLN2–HOPX
0.03
0.02
0.01
0.00
−0.01
−0.02
Module score
0
250
500
750
1000
Distance to PHF1+ neuron (μm)
B
Micro
0.06
0.04
0.02
0.00
−0.02
Module score
0
250
500
750
1000
Distance to PHF1+ neuron (μm)
C
Astro
0.02
0.01
0.00
−0.01
−0.02
Module score
0
250
500
750
1000
Distance to PHF1+ neuron (μm)
