## Supplementary material for "Neuronal and glial transcriptional states vary with proximity to tangle-bearing neurons in the human entorhinal cortex": Figure S3

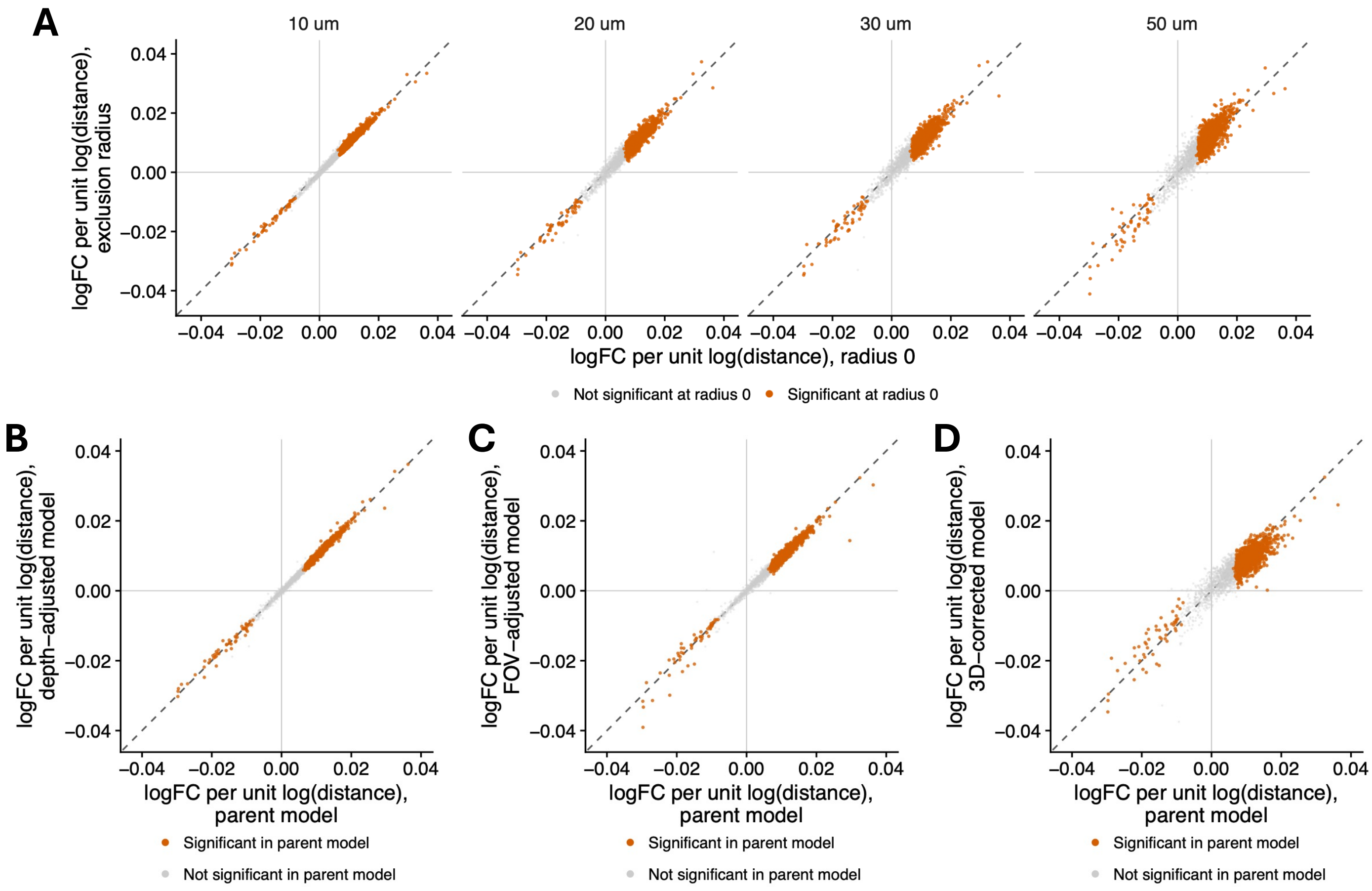
A
10 um
20 um
30 um
50 um
logFC per unit log(distance), exclusion radius
logFC per unit log(distance), radius 0
Not significant at radius 0
Significant at radius 0
B
logFC per unit log(distance), depth−adjusted model
logFC per unit log(distance), parent model
Significant in parent model
Not significant in parent model
C
logFC per unit log(distance), FOV−adjusted model
logFC per unit log(distance), parent model
Significant in parent model
Not significant in parent model
D
logFC per unit log(distance), 3D−corrected model
logFC per unit log(distance), parent model
Significant in parent model
Not significant in parent model
