## Supplementary material for "Neuronal and glial transcriptional states vary with proximity to tangle-bearing neurons in the human entorhinal cortex": Figure S4

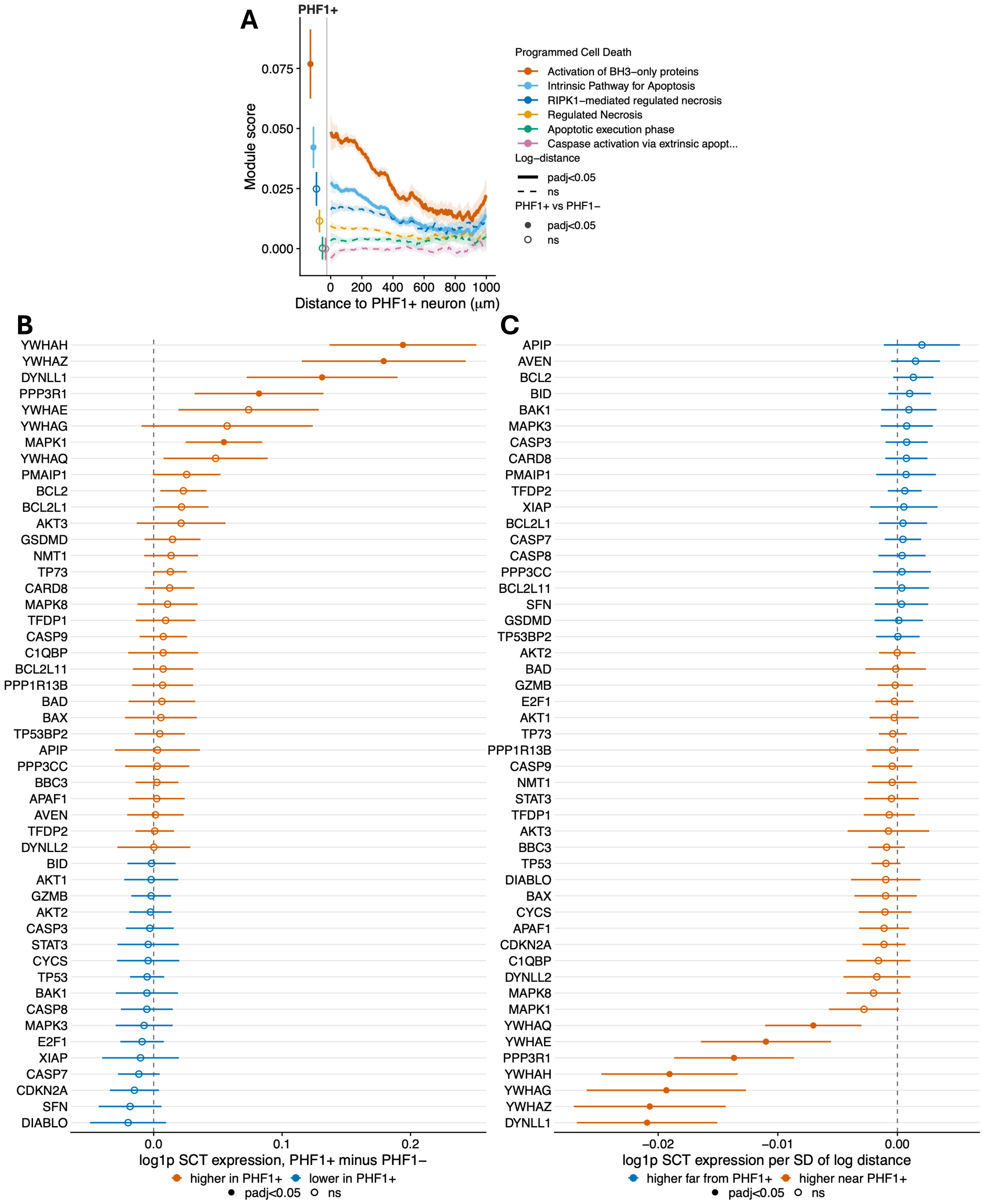
A
PHF1+
Module score
0.075
0.050
0.025
0.000
0
200
400
600
800
1000
Distance to PHF1+ neuron (μm)
Programmed Cell Death
Activation of BH3−only proteins
Intrinsic Pathway for Apoptosis
RIPK1−mediated regulated necrosis
Regulated Necrosis
Apoptotic execution phase
Caspase activation via extrinsic apopt...
Log−distance
padj<0.05
ns
PHF1+ vs PHF1−
padj<0.05
ns
B
YWHAH
YWHAZ
DYNLL1
PPP3R1
YWHAE
YWHAG
MAPK1
YWHAQ
PMAIP1
BCL2
BCL2L1
AKT3
GSDMD
NMT1
TP73
CARD8
MAPK8
TFDP1
CASP9
C1QBP
BCL2L11
PPP1R13B
BAD
BAX
TP53BP2
APIP
PPP3CC
BBC3
APAF1
AVEN
TFDP2
DYNLL2
BID
AKT1
GZMB
AKT2
CASP3
STAT3
CYCS
TP53
BAK1
CASP8
MAPK3
E2F1
XIAP
CASP7
CDKN2A
SFN
DIABLO
0.0
0.1
0.2
log1p SCT expression, PHF1+ minus PHF1−
higher in PHF1+
lower in PHF1+
padj<0.05
ns
C
APIP
AVEN
BCL2
BID
BAK1
MAPK3
CASP3
CARD8
PMAIP1
TFDP2
XIAP
BCL2L1
CASP7
CASP8
PPP3CC
BCL2L11
SFN
GSDMD
TP53BP2
AKT2
BAD
GZMB
E2F1
AKT1
TP73
PPP1R13B
CASP9
NMT1
STAT3
TFDP1
AKT3
BBC3
TP53
DIABLO
BAX
CYCS
APAF1
CDKN2A
C1QBP
DYNLL2
MAPK8
MAPK1
YWHAQ
YWHAE
PPP3R1
YWHAH
YWHAG
YWHAZ
DYNLL1
−0.02
−0.01
0.00
log1p SCT expression per SD of log distance
higher far from PHF1+
higher near PHF1+
padj<0.05
ns
