## Supplementary material for "Neuronal and glial transcriptional states vary with proximity to tangle-bearing neurons in the human entorhinal cortex": Figure S5

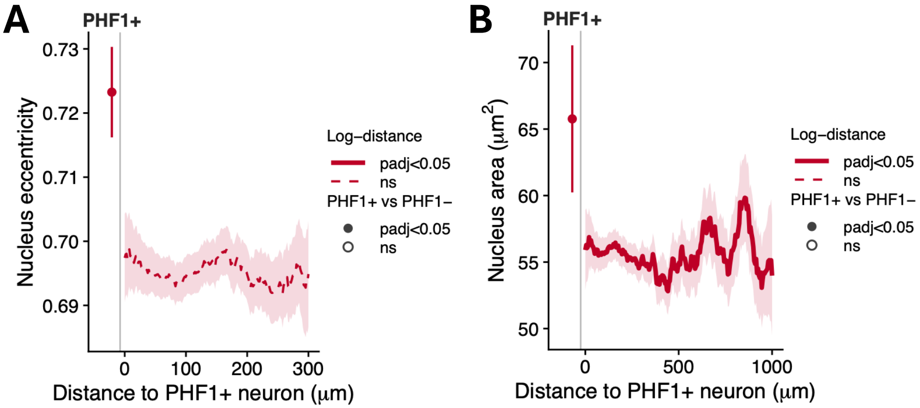

A
PHF1+
0.73
0.72
0.71
0.70
0.69
Nucleus eccentricity
0
100
200
300
Distance to PHF1+ neuron (μm)
Log−distance
padj<0.05
ns
PHF1+ vs PHF1−
padj<0.05
ns
B
PHF1+
70
65
60
55
50
Nucleus area (μm2)
0
500
1000
Distance to PHF1+ neuron (μm)
Log−distance
padj<0.05
ns
PHF1+ vs PHF1−
padj<0.05
ns
