## Supplementary material for "Neuronal and glial transcriptional states vary with proximity to tangle-bearing neurons in the human entorhinal cortex": Figure S6

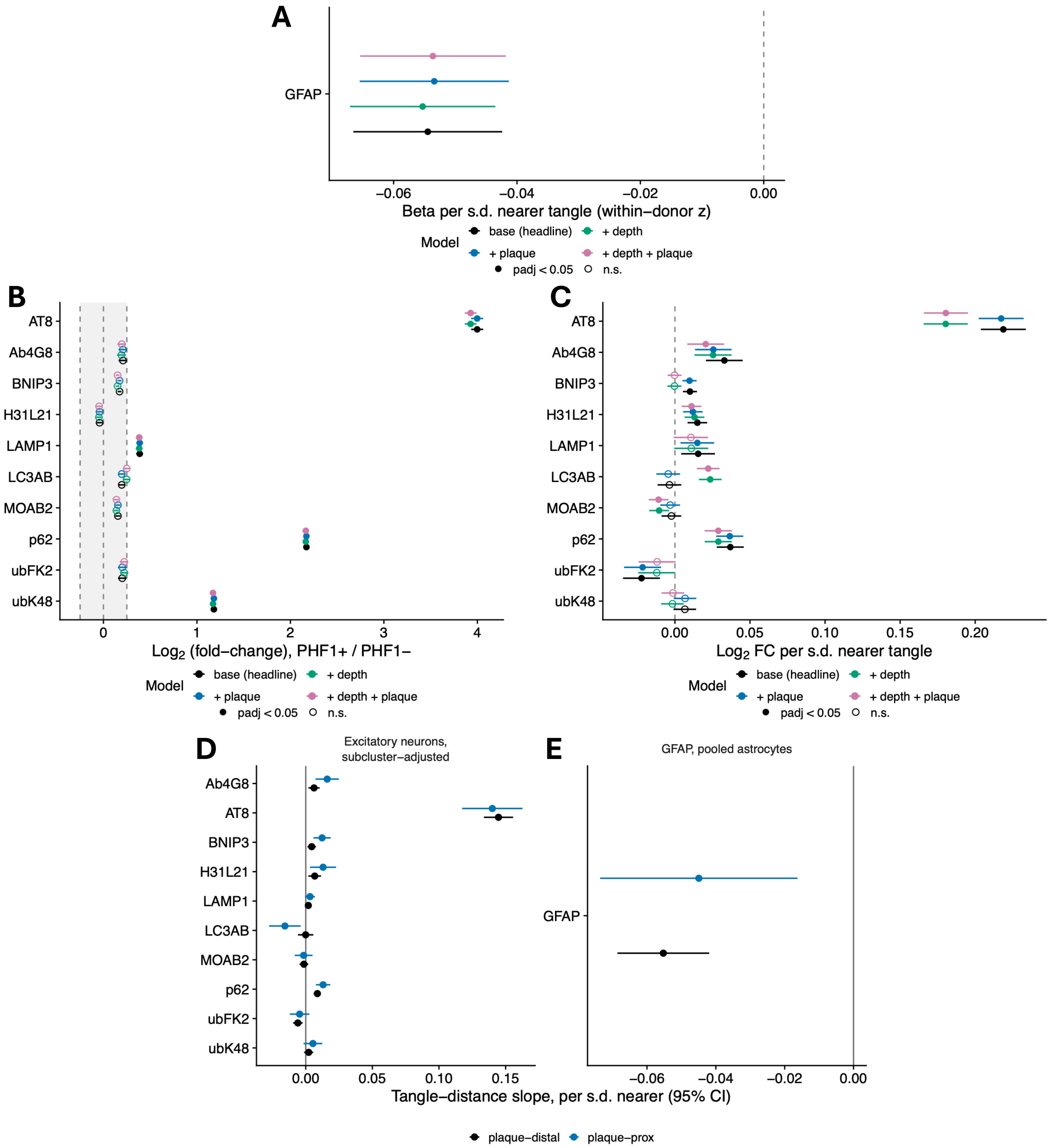
A
GFAP
−0.06
−0.04
−0.02
0.00
Beta per s.d. nearer tangle (within−donor z)
Model
base (headline)
+ depth
+ plaque
+ depth + plaque
padj < 0.05
n.s.
B
AT8
Ab4G8
BNIP3
H31L21
LAMP1
LC3AB
MOAB2
p62
ubFK2
ubK48
0
1
2
3
4
Log2 (fold−change), PHF1+ / PHF1−
C
0.00
0.05
0.10
0.15
0.20
Log2 FC per s.d. nearer tangle
D
Excitatory neurons, subcluster−adjusted
0.00
0.05
0.10
0.15
E
GFAP, pooled astrocytes
−0.06
−0.04
−0.02
0.00
Tangle−distance slope, per s.d. nearer (95% CI)
plaque−distal
plaque−prox
