## Supplementary material for "Neuronal and glial transcriptional states vary with proximity to tangle-bearing neurons in the human entorhinal cortex": Figure S7

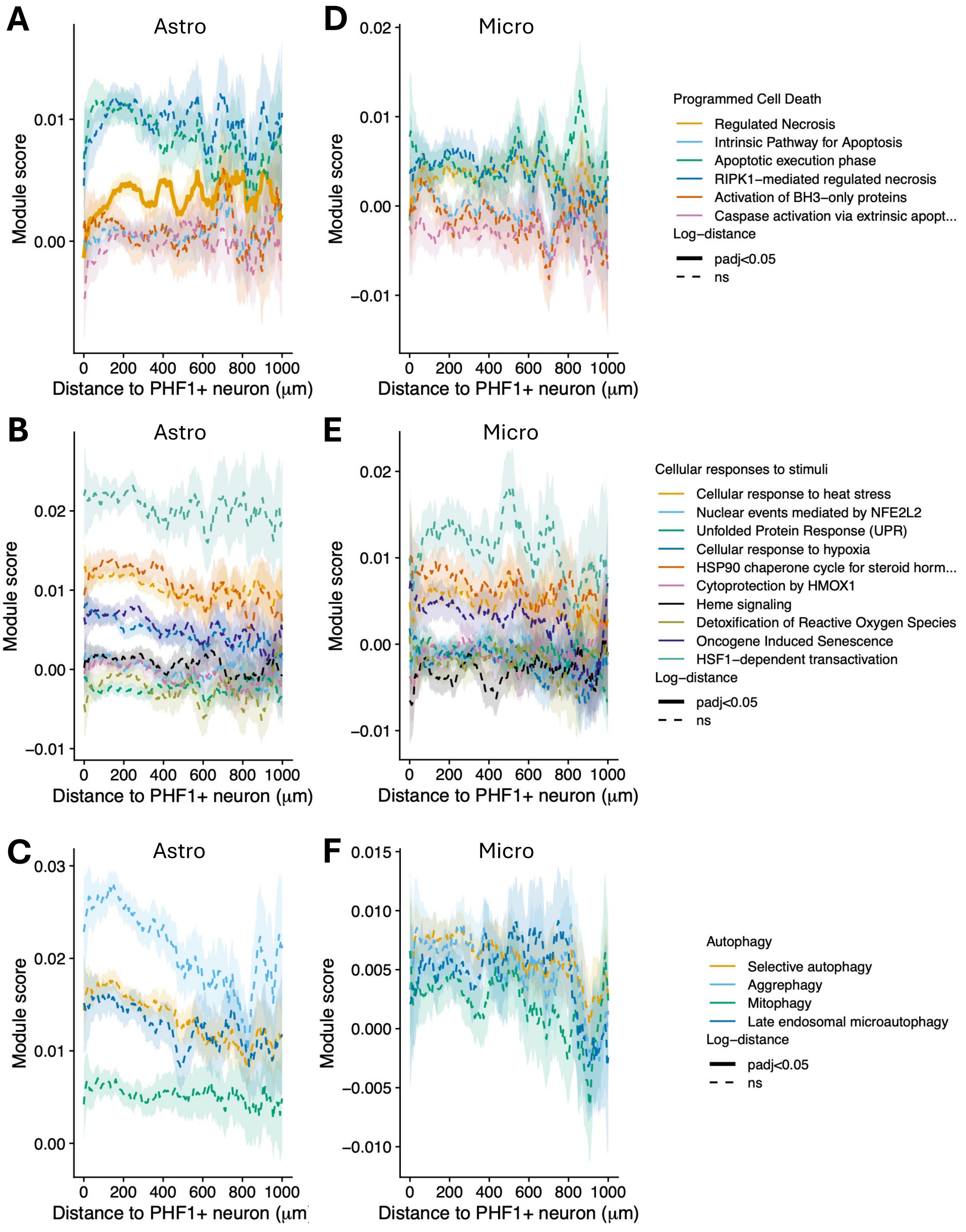

A
Astro
D
Micro
Programmed Cell Death
Regulated Necrosis
Intrinsic Pathway for Apoptosis
Apoptotic execution phase
RIPK1−mediated regulated necrosis
Activation of BH3−only proteins
Caspase activation via extrinsic apopt...
Log−distance
padj<0.05
ns
B
Astro
E
Micro
Cellular responses to stimuli
Cellular response to heat stress
Nuclear events mediated by NFE2L2
Unfolded Protein Response (UPR)
Cellular response to hypoxia
HSP90 chaperone cycle for steroid horm...
Cytoprotection by HMOX1
Heme signaling
Detoxification of Reactive Oxygen Species
Oncogene Induced Senescence
HSF1−dependent transactivation
Log−distance
padj<0.05
ns
C
Astro
F
Micro
Autophagy
Selective autophagy
Aggrephagy
Mitophagy
Late endosomal microautophagy
Log−distance
padj<0.05
ns
Module score
Distance to PHF1+ neuron (μm)
